# Re-evaluating frontopolar and temporoparietal contributions to detection and discrimination confidence

**DOI:** 10.1101/2022.08.23.503975

**Authors:** Matan Mazor, Chudi Gong, Stephen M. Fleming

**Author notes:** **Author Note:** This research was funded in whole, or in part, by the Wellcome Trust [Grant number 203147/Z/16/]. For the purpose of open access, the author has applied a CC BY public copyright licence to any Author Accepted Manuscript version arising from this submission.

## Abstract

Previously, we identified a subset of regions where the relation between decision confidence and univariate fMRI activity was quadratic, with stronger activation for both high and low compared to intermediate levels of confidence. We further showed that, in a subset of these regions, this quadratic modulation appeared only for confidence in detection decisions about the presence or absence of a stimulus, and not for confidence in discrimination decisions about stimulus identity (Mazor, Friston & Fleming, 2021). Here, in a pre-registered follow-up experiment, we sought to replicate our original findings and identify the origins of putative detection-specific confidence signals by introducing a novel asymmetric-discrimination condition: a discrimination task with the signal-detection properties of a detection task. This task required discriminating two alternatives (two different grating tilts) but was engineered such that the distribution of perceptual evidence was asymmetric, just as in yes/no detection. We successfully replicated the quadratic modulation of subjective confidence in prefrontal, parietal and temporal cortices. However, in contrast to our original report, this quadratic effect was similar in detection and discrimination responses, but stronger in the novel asymmetric-discrimination condition. We interpret our findings as weighing against the detection-specificity of confidence signatures and speculate about possible alternative origins of a quadratic modulation of decision confidence.

## Introduction

Adult humans are able not only to evaluate what they see and don’t see, but also how confident they are in these percepts (Mamassian, 2016). Investigations into the neural basis of metacognition reveal a network of brain regions where activation scales with perceptual confidence (for a coordinate-based meta-analysis, see Vaccaro & Fleming, 2018). However, a majority of previous computational modelling and neuroimaging studies of perceptual confidence have focused on understanding confidence in discrimination decisions (e.g., *was it a bird or a plane?*). In contrast, the computations and neural substrates supporting perceptual confidence for detection decisions (e.g., *was there anything there at all?*) remain largely uncharted territory. Mapping that territory is of considerable interest, both due to the conceptual overlap between (detection) confidence and perceptual awareness, and also because detection may invoke distinct computational demands that are not required in discrimination (Triesman & Williams, 1984; Ko & Lau, 2012; Fleming, 2020).

In a previous study (Mazor et al., 2020) we compared the parametric effect of subjective decision confidence on brain activation in two perceptual decision-making tasks: a discrimination task (*was the grating tilted clockwise or anticlockwise?*) and a detection task (*was there any grating present at all?*). Replicating previous findings (Bang & Fleming, 2018; Morales et al., 2018; Vaccaro & Fleming, 2018), we observed a linear effect of confidence in a set of pre-defined regions of interest, with high confidence levels associated with a stronger (ventromedial prefrontal cortex, vmPFC; precuneus; ventral striatum) or weaker (posterior medial frontal cortex, pMFC) signals, across both tasks and responses. Exploratory analysis additionally revealed a widespread positive quadratic effect of confidence, with stronger signals associated with using the extreme ends of the confidence scale. In the right frontopolar cortex, right superior temporal sulcus (STS) and right pre-supplementary motor area (pre-SMA) this quadratic effect was stronger for the detection task, where participants decided whether a grating was present or absent. Additionally, in the right temporoparietal junction (rTPJ), a linear effect of confidence was stronger following judgments about target absence compared with judgments about target presence.

Signal-detection based computational simulations suggested that a quadratic activation profile may reflect the unequal variance nature of detection tasks (see Fig. 1). In detection, the variance associated with perceiving a signal is higher than the variance associated with perceiving the absence of a signal (Wickens, 2001; Kellij et al., 2021). This unequal variance evidence structure can then produce a quadratic activation pattern in brain regions that are involved in dynamically updating a decision criterion or in representing the likelihood ratio between the two stimulus classes (Mazor et al., 2020). An alternative interpretation of our previous results is that distinct metacognitive processes are selectively invoked for decisions and confidence formation about presence and absence, but not in confidence about stimulus identity. For example, brain regions that selectively encode stimulus visibility (Mazor, Dijkstra & Fleming, 2022), ones that correspond to higher-order nodes in a hierarchical model of perceptual states (Fleming, 2020), or ones that are implicated in counterfactual thinking and attention monitoring (Mazor, 2021) may show differential modulation of confidence in detection and discrimination decisions.

**Fig. 1:**
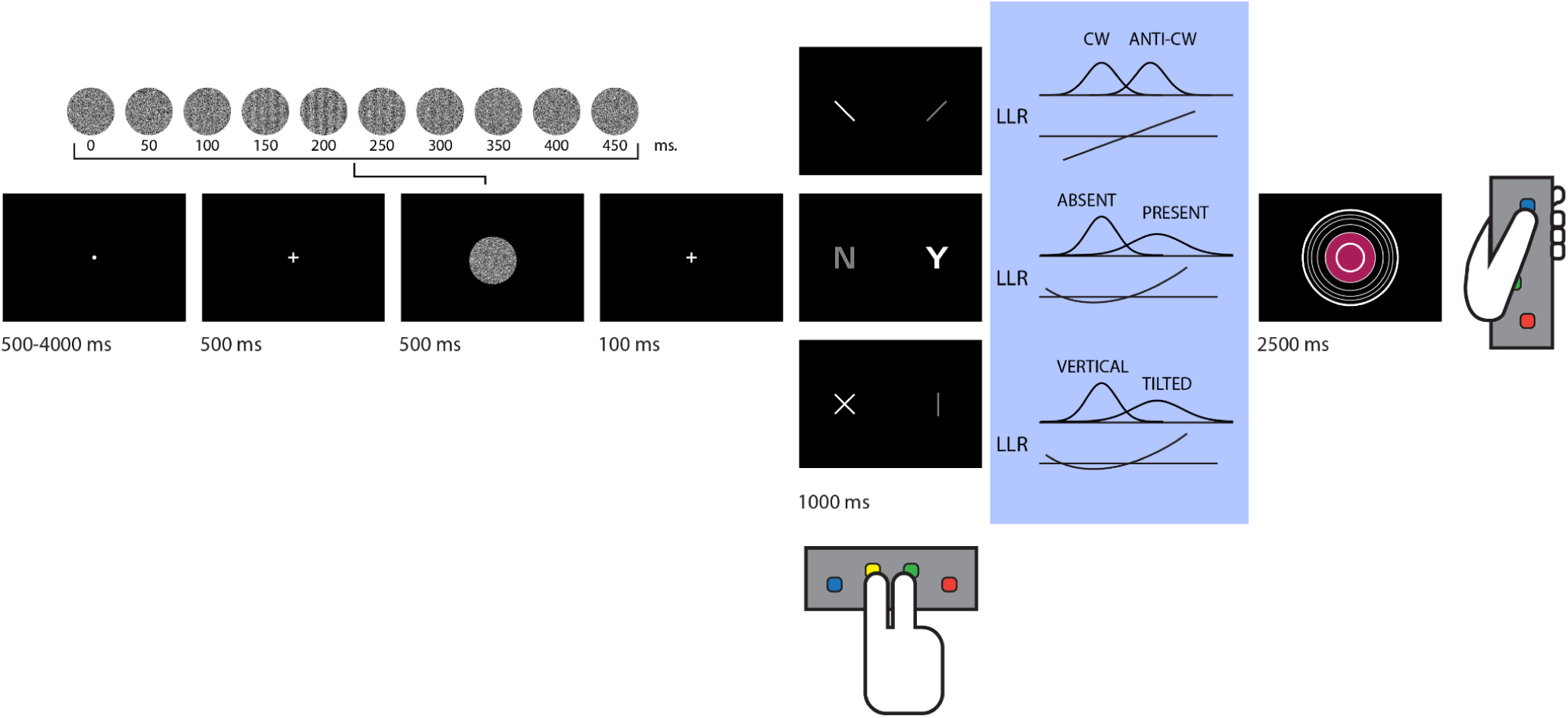
Experimental design. Stimuli consisted of dynamic random patterns of grayscale values. In all trials except for detection ‘target absent’ trials, a grating emerged from and disappeared back into the noise. Participants used the index and middle fingers of their right hand to indicate whether a grating was tilted clockwise or anticlockwise (discrimination), whether it was present or absent (detection), or whether it was vertical or tilted (tilt recognition). They then reported their level of confidence on a 6-point scale by controlling the size of a coloured circle with their left thumb. In blue: inter-trial variability was similar for the two stimulus categories in discrimination, whereas in detection and tilt recognition it was higher for one category over the other. Log Likelihood Ratio (LLR) is a linear function of the perceptual sample in discrimination, but this relation is quadratic in detection and tilt recognition.

The design of our previous study did not allow us to decide between these alternative accounts. Here, we introduce a third hybrid condition to our experimental design: a discrimination task with the distributional properties of a detection task (*tilt recognition*; following Denison et al., 2018). This task requires subjects to report whether a grating is tilted or vertical: a discrimination judgment between two stimulus classes. However, because tilted gratings can appear in various orientations while vertical gratings are fixed, the distribution of perceptual evidence is of higher variance in the former, mimicking the variance asymmetry of a (yes/no) detection task. The two possible explanations of our previous findings (sensitivity of confidence encoding to variance structure versus a specific representation of presence and absence) thus make different predictions for this third condition. An unequal-variance account predicts qualitatively similar neural confidence effects to those observed in detection, as the two tasks share a similar distributional structure. Conversely, a presence/absence asymmetry account predicts confidence effects that are qualitatively similar to those observed in discrimination, as the tilt-recognition task no longer requires inference about stimulus presence vs. absence.

To anticipate our results, behavioural analysis confirmed that the tilt-recognition task induced detection-like unequal-variance effects on subjects’ confidence ratings – confirming that it created an asymmetric discrimination task, as intended. A mass univariate analysis of fMRI data replicated linear and quadratic effects of confidence in pre-specified regions of interest. However, unlike in our previous study, here the quadratic effects of confidence were now similar in the detection and discrimination tasks, and were instead stronger in the novel asymmetric-discrimination condition, which was also the condition with the most pronounced behavioural signatures of unequal variance. Furthermore, and in contrast to what we observed in Mazor et al. (2020), in the rTPJ, a negative linear modulation of decision confidence was similar in detection ‘yes’ and ‘no’ responses. Representational similarity analysis indicated that differences in multivariate activity patterns between high- and low-confidence trials were mostly task-invariant. We conclude with a discussion of how our previous conclusions should be revised in light of these new results.

## Results

In a pre-registered design (pre-registered protocol folder: github.com/matanmazor/unequalVarianceDiscrimination/tree/main/experiment/protocolFolder/pr otocolFolder), a total of 46 participants performed three perceptual decision-making tasks while being scanned in a 3T MRI scanner: an orientation discrimination task (‘was the grating tilted clockwise or anticlockwise?‘), a detection task (‘was any grating presented at all?’), and a tilt-recognition task (’was the grating tilted or vertical?’; see Figure 1). Tasks were performed in separate blocks each comprising 26 trials. At the end of each trial, participants rated their confidence in the accuracy of their decision on a 6-point scale. We adjusted the difficulty of the three tasks in a preceding behavioral session to achieve similar performance of around 70% accuracy. 15-18 blocks were presented in 5-6 scanner runs.

### Behavioural Results

35 participants met our pre-registered inclusion criteria (see Methods). Task performance was similar for discrimination (76% accuracy), detection (78% accuracy), and tilt recognition (77% accuracy). Repeated measures analysis of variance revealed no difference in response accuracy between the three tasks (*F(2,68)=1.17, p=.32, BF_01_*=3.43; see Fig. 2A). The probability of responding ‘clockwise’ in the discrimination task was 51% and not significantly different from 0.5 (*t(34)=0.58, p=.57, d=0.10*). In contrast, participants were more likely to respond ‘no’ than ‘yes’ in the detection task (54% of all responses, *t(34)=3.70,p<.001,d=0.63*), and ‘vertical’ than ‘tilted’ in the tilt recognition task (57% of all responses, *t(34)=6.67,p<.001,d=1.13*). This is consistent with an optimal setting of a decision criterion in an unequal-variance setting (Rahnev, 2021).

**Fig. 2:**
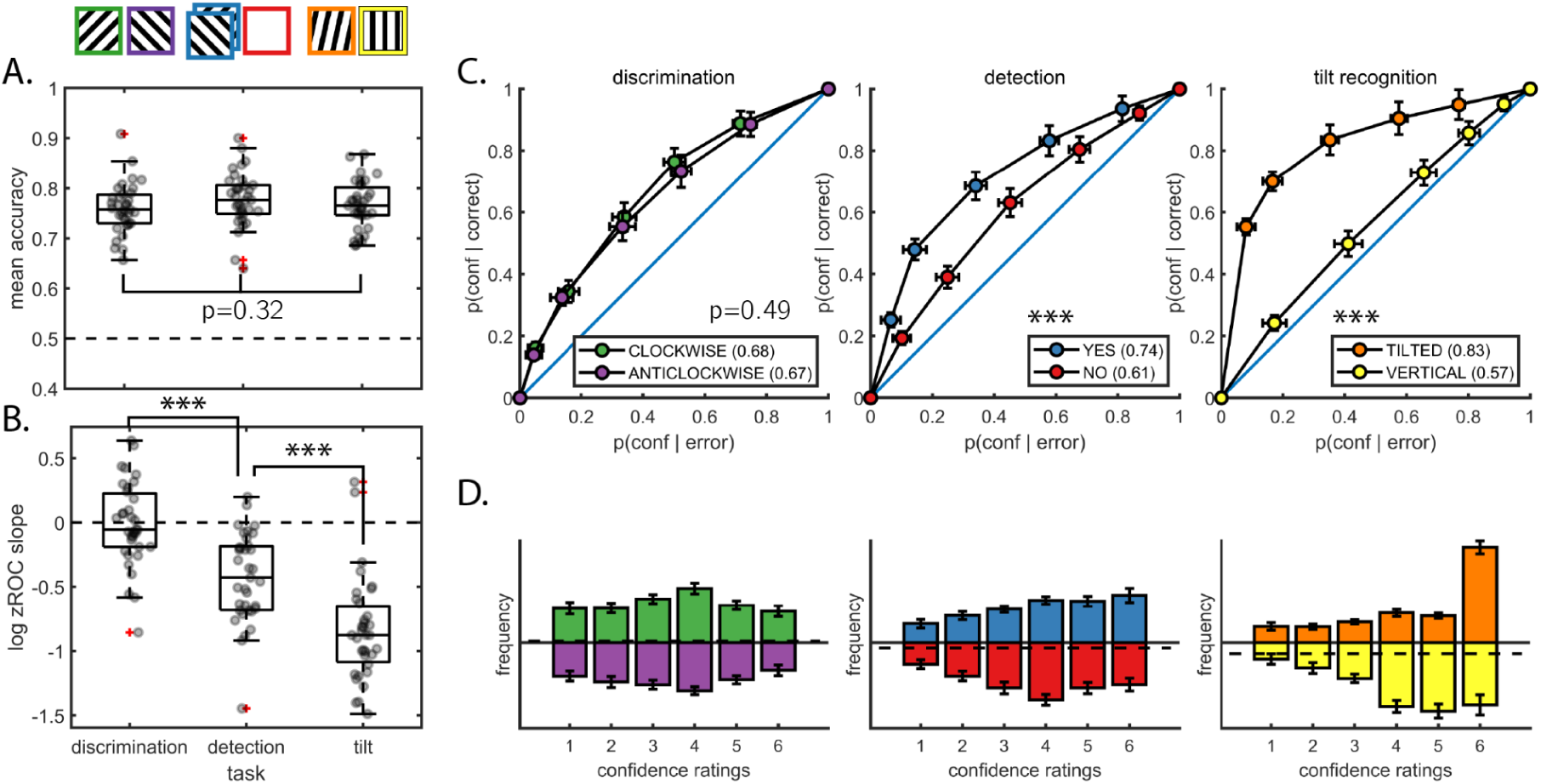
Behavioural results. A: response accuracy was similar for the three tasks. B: the log zROC slope was not different from 0 in discrimination, indicating similar variability in the representation of clockwise and anticlockwise stimuli. In detection, this quantity was significantly negative, indicating higher variability in the representation of signal. In tilt recognition, this quantity was even more negative, indicating higher variability in the representation of tilted stimuli. C. metacognitive sensitivity, quantified as the area under the response-conditional type-II ROC curve, was significantly higher for both ‘yes’ and ‘tilted’ responses compared to ‘no’ and ‘vertical’ responses, respectively. We observed no significant difference in metacognitive sensitivity between discrimination ‘clockwise’ and ‘anticlockwise’ responses. D. distributions of confidence ratings (on a 1-6 scale) for the three tasks and six responses.

Similar to what we had observed in Mazor at al. (2020), participants were equally confident in reporting both clockwise (mean confidence on a 1-6 scale: *3.51*) and anticlockwise tilt (mean confidence: *3.44*) in the discrimination task (*t(34)=0.80,p=.43,d=0.14*). However, unlike in our previous study, here a numerical difference in mean confidence between detection ‘yes’ (*3.95*) and ‘no’ (*3.80*) responses was not significant (*t(34)=1.62,p=.12, d=0.27*). Finally, in the tilt recognition task participants were significantly more confident in reporting a tilted grating (*4.60*) than compared to a vertical grating (*4.19*; *t(34)=3.71, p<.001,d=0.63;* see Fig. 2D).

Confidence ratings are typically aligned with objective accuracy, such that participants are more confident on average when they are correct, compared to when they are wrong. Previous studies found that this alignment, commonly referred to as *metacognitive sensitivity*, is reduced for decisions about target absence compared to decisions about target presence (Kellij et al., 2021; Mazor et al., 2020, 2021; Meuwese et al., 2014). Here also, metacognitive sensitivity (quantified as the area under the response-conditional type-II ROC curve), was significantly higher for detection ‘yes’ compared to ‘no’ responses (*t(34)=6.41,p<.001,d=1.1;* see Fig. 2C). Similarly, in the tilt recognition task metacognitive sensitivity was higher for ‘tilted’ compared to ‘vertical’ responses (*t(34)=9.55, p<.001,d=1.61*). No difference in metacognitive sensitivity was observed between discrimination clockwise and anticlockwise responses (*t(34)=0.70, p=.49,d=0.12*).

In an unequal-variance signal detection setting, metacognitive sensitivity is expected to be higher for classifying a stimulus as belonging to the high-variance compared to the low-variance stimulus class. Our tilt recognition task is an example of such a setting: the ‘vertical’ class had low variance (all stimuli were vertical), and the ‘tilted’ class had high variance (some stimuli were more tilted than others). As expected, the ratio between the standard deviations of the two stimulus categories (measured as the geometric mean of type-1 zROC slopes), was *0.55* and significantly lower than 1, indicating higher variability in the representation of tilted stimuli (a t-test performed on log-slopes against 0: *t(34)=-12.50, p<.001, d=2.11*; see Fig. 2B). Similarly, this ratio was *0.74* for the detection task, indicating higher variability in the encoding of target presence (*t(34)=-7.27,p<.001,d=1.23*). In contrast, the ratio was *0.99* for the discrimination task and statistically indistinguishable from 1, indicating similar variability in the encoding of clockwise and anticlockwise stimuli (*t(34)=-0.26,p=0.79,d=0.04*).

### Imaging results

Whole brain results for all pre-registered contrasts are available on NeuroVault (https://identifiers.org/neurovault.collection:12352). Anonymized imaging data from all included subjects is available on OpenNeuro (https://openneuro.org/datasets/ds004081/versions/1.0.0). We pre-registered a plan to evaluate the parametric modulation of confidence both directly from brain activations, as well as indirectly from beta coefficients of a design matrix where confidence is specified as a categorial variable. This two-step solution controls for metacognitive biases in that all confidence levels equally contribute to modulation estimates, regardless of their frequency in the data. Results from the categorical design matrices mostly agreed with those from the parametric modulation analysis. We therefore report the parametric modulation results, and mention when the categorical design matrix provided conflicting results. Full results from both approaches are presented in the supplementary.

#### Linear and Quadratic effects of confidence

Our primary (Quadratic Confidence) design matrix included parametric modulators for linear and quadratic effects of confidence. Among our pre-specified regions of interest, the medial frontopolar (FPm) and ventromedial prefrontal cortex (vmPFC) ROIs showed a positive linear effect of confidence (FPm: *t(34)=3.45, p<.001, d=0.58*, vmPFC: *t(34)=4.53, p<.001, d=0.77*). A whole brain contrast revealed a positive modulation of confidence in the bilateral precuneus, claustrum, and ventral striatum. Conversely, the right temporoparietal junction showed a negative linear modulation of confidence, similar to what we observed in our previous study (*t(34)=-3.35,p<.001,d=0.57*). Quadratic polynomials fitted to beta values from the categorical design matrices revealed a negative linear modulation of confidence also in the right superior temporal sulcus (*t(34)=2.47, p<0.05, d=0.42*), and a whole-brain analysis revealed a negative linear effect of confidence in the posterior middle frontal cortex (pMFC).

Consistent with what we had observed in our previous study, a positive quadratic effect of confidence was robustly observed in a number of regions. Among our pre-specified ROIs, this effect was significant in lateral frontopolar cortex (FPl; *t(34)=2.93,p<.01, d=0.50*), Brodmann area 46 (BA46; *t(34)=4.43,p<.001, d=0.75*), rTPJ (*t(34)=4.89,p<.001, d=0.83*), right superior temporal sulcus (rSTS; *t(34)=3.62,p<.001, d=0.61*), and pre-supplementary motor area (preSMA; *t(34)=5.00, p<0.001, d=0.85*). Whole-brain analysis revealed a quadratic effect of confidence also in dorsolateral prefrontal and orbitofrontal cortex, anterior insula, precuneus, posterior cingulate, and in the cerebellum.

#### Task-specific activations

We next asked whether brain activation differed between the three tasks, collapsed across responses and confidence levels. Repeated measures analyses of variance failed to find a main effect of task in any of our 7 pre-registered ROIs (all p’s>.33; see S3). Outside these regions, whole-brain analysis (*p<*0.05, corrected for family-wise error at the cluster level) revealed that activations in bilateral premotor cortex were sensitive to task identity. This is consistent with the successful encoding of semantic meaning of motor actions from associative motor cortex (Aberbach et al., 2021).

#### Task- and response-specific confidence modulations

We next asked whether confidence-related brain activation differed between the three tasks. A linear modulation of confidence was similar for the three tasks: repeated measures ANOVAs revealed no effect of task in any of our ROIs (all p’s>.30; see S4 and Fig. 3, first row), and whole-brain analysis revealed no differences outside these pre-specified regions.

**Fig. 3:**
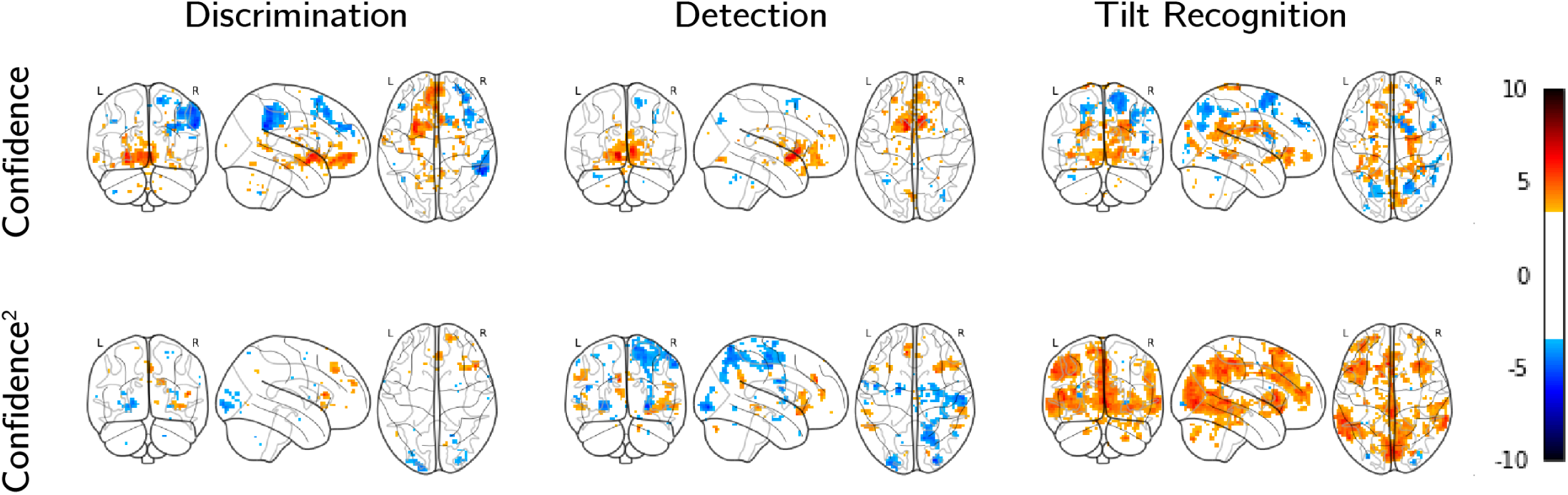
Linear (first row) and quadratic (second row) effects of confidence for the three tasks. *P* < 0.001, uncorrected for multiple comparison.

Our main hypothesis however was that a quadratic modulation of confidence should be more pronounced in some tasks than in others – indicating presence/absence or unequal-variance-related asymmetries. Within our ROIs, this was the case in BA46 (*F(2,34)=4.47,p<.05*), rSTS (*F(2,34)=3.80,p<.05*), and marginally in rTPJ (*F(2,34)=2.1,p=0.08*). The same regions showed a marginal effect when subjecting betas from the categorical design matrix to a group level ANOVA (*p=0.06*, *p=0.08* and *p=0.06* for BA46, rSTS, and rTPJ, respectively). Whole-brain analysis further revealed differences in a quadratic modulation of confidence across tasks in the insula (*p*<0.05, cluster-corrected).

Our next set of pre-registered tests was designed to pinpoint the origins of this interaction of the quadratic expansion of confidence with task. First, we attempted to replicate our finding from study 1 of a stronger quadratic modulation of confidence in detection compared to discrimination. Contrary to our prediction, we found no significant differences in modulation in any of our ROIs (all p’s>0.17, see S6). To determine whether this absence of a significant result should also be taken as positive evidence against a difference between detection and discrimination, we subjected our data to a Bayesian t-test (Rouder et al., 2009). In the FPm ROI, we obtained moderate evidence *against* a difference between detection and discrimination (*BF_01_=3.71*). Similarly, we obtained moderate evidence against a difference between detection and discrimination in the pre-SMA (*BF_01_=5.43*). Analysing beta values from the categorical design matrix, we obtained moderate evidence for the null hypothesis of no difference also in FPl (*BF_01_=3.04*) and rTPJ (*BF_01_=5.48*). Bayes factors for all other ROIs in which we observed an effect in study 1 were within the interval [⅓,3], indicating no clear evidence for or against an effect (see S6).

In contrast to detection, where quadratic confidence profiles were similar to discrimination, the quadratic effect of confidence was significantly stronger in the tilt recognition relative to the discrimination task in BA46 (*t(34)=2.23, p<.05*), rTPJ (*t(34)=2.18, p<.05*), rSTS (*t(34)=2.86, p<0.01*), and marginally in preSMA (*t(34)=1.78,p=0.08*).

In our original study (Mazor et al., 2020), a cluster in the right temporoparietal junction showed a negative linear effect of confidence, which was significantly more negative in detection ‘no’ compared to ‘yes’ responses. In the current study, activation in this region again showed a negative linear, as well as a positive quadratic effect of confidence. However, an interaction between confidence and detection response was not significant (*t(34)=0.72,p=.47*). A Bayesian t test provided moderate evidence against a difference between detection ‘yes’ and ‘no’ responses in this region (*BF_01_=4.33*). No other region of interest showed an interaction of confidence with detection response (all p’s>0.23, see S8). Similarly, none of our ROIs showed a significant interaction of confidence with response in the tilt recognition task (all p’s>0.11; see S9). A whole-brain analysis revealed a cluster in the left lingual gyrus (MNI coordinates [-30,-64, −8]) in which a linear modulation of confidence was stronger in ‘tilt’ compared to ‘vertical’ responses (*p*<0.05, cluster-corrected)..

#### Multivariate analysis

To further investigate the relationships between spatial activation patterns across tasks, responses, and confidence levels, we next turned to Representational Similarity Analysis (RSA; Kriegeskorte et al., 2008). Specifically, we asked which regions represented task, irrespective of confidence; which represented confidence, irrespective of task; and which represented confidence in a task-dependent manner. We pre-registered eight representational dissimilarity matrices (RDMs), each specifying a theory-based prediction regarding which trials should be encoded similarly or dissimilarly based on task, response and reported confidence (see Fig. 4) and compared them against empirical dissimilarity matrices extracted from our pre-registered ROIs.

**Fig. 4:**
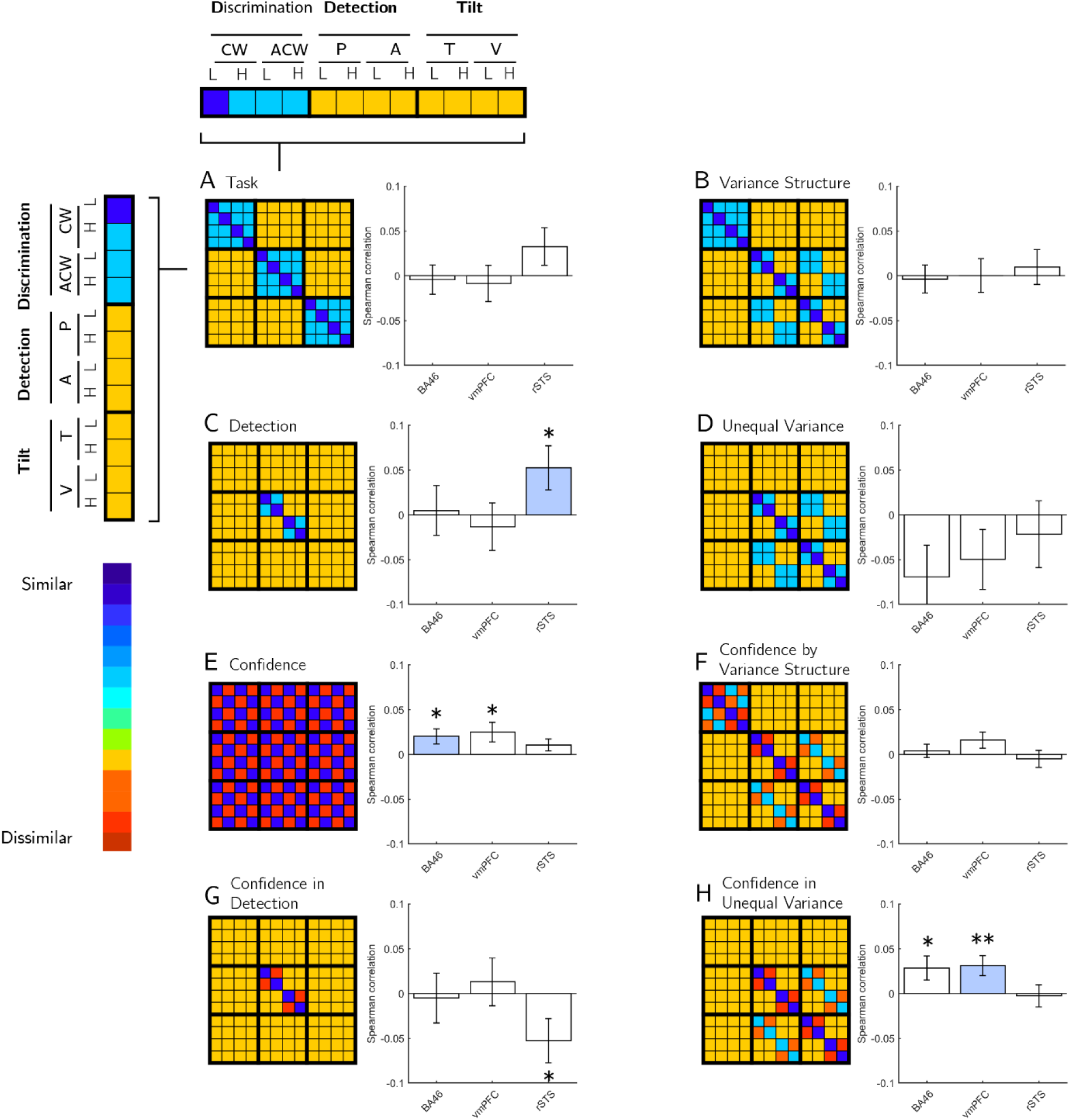
The eight a-priori Representational Dissimilarity Matrices (RDMs) used for Representational Similarity Analysis, and their corresponding Spearman correlations with empirical RDMs from the three ROIs. For each ROI, we marked the RDM that produced the most robust correlation (in standardized effect sizes) in blue.

As a first step, and in order to verify that empirical RDMs hold reliable condition-specific multivariate activation patterns, we compared on- and off-diagonal entries in the ranked RDMs. If condition-specific information is encoded in RDMs, off-diagonal distances should be higher than on-diagonal ones (Ritchie et all., 2017). This was the case in FPm (t(34)=3.29, p<0.01), BA46 (t(34)=2.87, p<0.01), vmPFC (t(34)=2.06, p<0.05) and rSTS (t(34)=2.17, p<0.05), but not in FPl, rTPJ and pre-SMA (all p’s>0.17). In the FPm ROI, subject specific RDMs were not predicted by the averaged RDM of other subjects (p=0.38), reflecting poor multivariate signal in this region (Nili et al., 2014). We therefore restricted the multivariate analysis to the three regions with reliable multivariate activation patterns: BA46, vmPFC, and rSTS. In all following analyses, on-diagonal entries are ignored, in order not to artificially inflate correlation measures (Ritchie et al., 2017).

In the rSTS ROI, multivariate activation patterns were most consistent with encoding of detection responses (target present or absent; RDM C, see Fig. 4C; p<0.05). A negative correlation with RDM G (detection confidence) is driven by the perfect negative correlation between RDMs C and G, when excluding the diagonal entries.

In contrast, in the prefrontal BA46 and vmPFC ROIs, multivariate brain activation patterns were most consistent with task-invariant confidence encoding (RDM E, see Fig. 4E; p’s<0.05), and with confidence encoding that is specific to an unequal-variance setting (RDM H, see Fig. 4H, p’s<0.05). Spearman correlations with RDMs E and H were not significantly different from each other (p’s>0.54). This is in line with our results from our previous study (Mazor et al., 2020), where cross-classification analysis revealed no evidence for task-specificity in multivariate confidence representations (see full pre-registered analyses https://osf.io/y3ftk/, Section “Task-specific and task-invariant confidence representation”).

To further explore the sensitivity of confidence encoding to variance structure we subjected the empirical RDMs to a multiple regression analysis in which candidate RDMs competed to explain the variance in the empirical RDM. First, we decomposed RDM E into 18 constituent sub-RDMs. Each such RDM represented the similarity between confidence encoding in two tasks, or within a single task (see Fig. 5A). We focused on specific sub-RDMs for which a variance structure account made unique predictions. In particular, a variance structure account predicts a response-specific similarity between confidence in detection and in tilt recognition (see Fig. 5B), but a presence-absence account predicts a similar response-invariant encoding of confidence in tilt-recognition and in discrimination (see Fig. 5C). To test this prediction, we compared the weighted coefficient combinations for these two predictions.

**Fig. 5:**
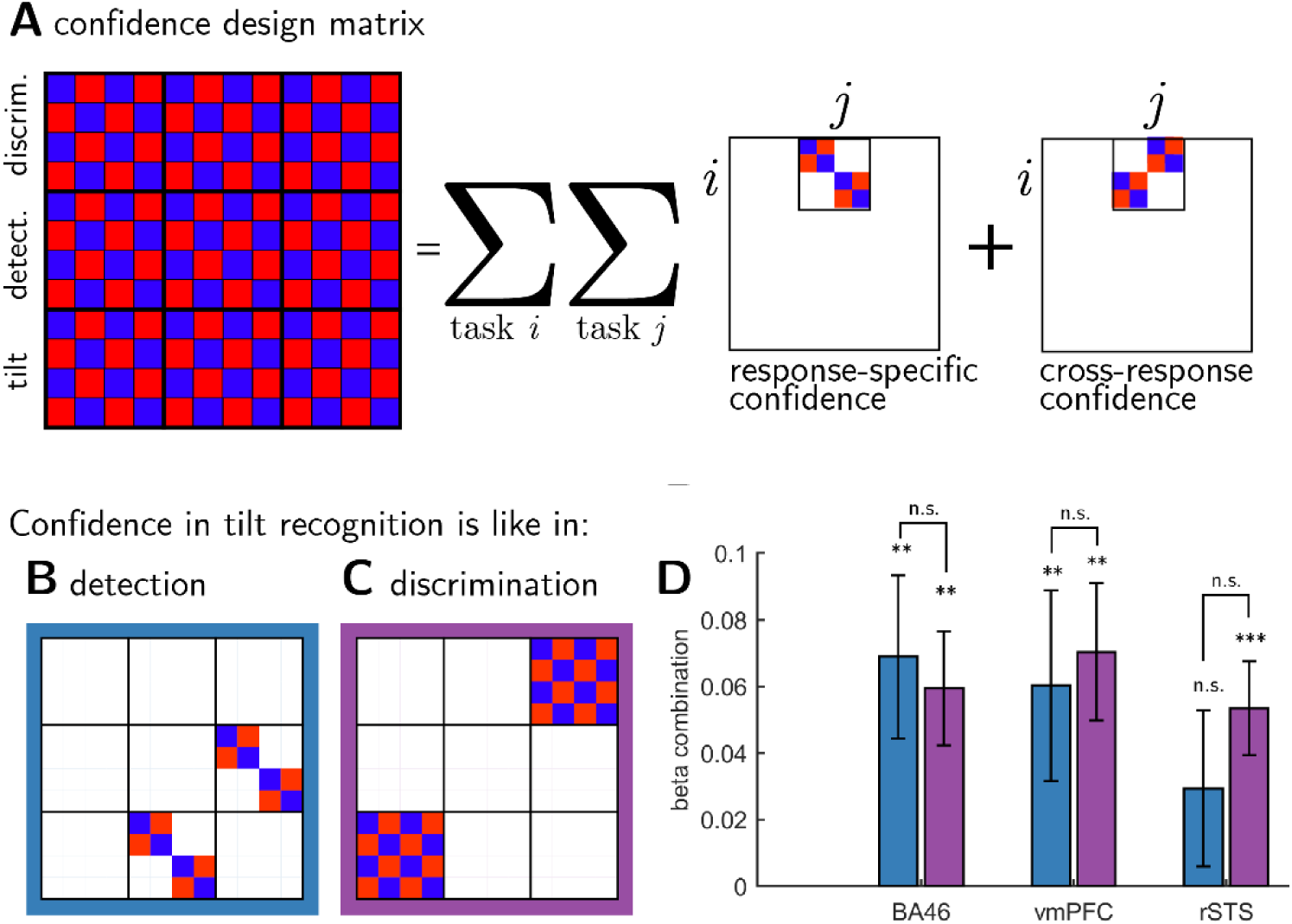
Multiple regression analysis. RDM E from Figure 4 was broken down into 18 constituent RDMs, which were then used to predict empirical RDMs in the seven ROIs (panel A). We then used beta coefficients from this multiple regression analysis to produce two beta combinations encoding our two hypotheses about differences between tasks (B and C): The first corresponds to similarity in confidence encoding between the tilt-recognition and the detection tasks, and the second to similarity in confidence-encoding between the tilt-recognition and discrimination tasks. We found no significant differences between these two weighted contrasts (panel D), supporting a task-invariant account of confidence encoding in these regions.

In the two prefrontal regions, vmPFC and BA46, multivariate activation patterns in the tilt recognition task were similar both to multivariate activation patterns in the discrimination task (vmPFC: t(34)=3.41, p<0.01; BA46: t(34)=3.48, p=0.001), and to multivariate activation patterns in the detection task (vmPFC: t(34)=2.10, p<0.05; BA46: t(34)=2.81, p<0.01). Consistent with our RSA results, this finding is in line with a task-invariant multivariate representation of confidence in these regions.

In contrast, rSTS showed similar confidence encoding in the tilt-recognition and discrimination tasks (t(34)=3.79, p<0.001), but distinct confidence encodings in tilt-recognition and detection tasks (t(34)=1.25, p=.22). Still, we observed no significant difference between these correlations (t(34)=0.84, p=0.41; see Fig. 5D), providing only indirect support for the hypothesis that multivariate confidence encoding in rSTS was different for detection and discrimination.

## Discussion

In a previous study we identified distinct neural contributions to confidence in perceptual detection that were not observed in a performance-matched discrimination task. In this pre-registered follow-up study, we set out to replicate this finding, and to further characterise the computational origins of a quadratic modulation of confidence. In what follows, we summarise our findings, discuss how our previous conclusions should be revised in light of this new data, and unpack what this may mean for our understanding of a quadratic effect of decision confidence in association cortex.

### A quadratic modulation of confidence is not distinct to perceptual detection

As in Mazor et al. (2020), here too we observed a widespread quadratic modulation of subjective confidence in prefrontal and parietal association cortex. In our previous study, this effect was stronger in a perceptual detection task, and in some frontopolar regions was not observed at all in a performance-matched discrimination task. In contrast, in the present study this effect was no longer specific to detection, and was instead observed in both detection and discrimination tasks.

There are a number of potential reasons for this difference between our results here and those reported in our previous study; for example, in our previous study stimuli were presented briefly, whereas here we opted for a dynamic display mode (see Methods). Alternatively, the inclusion of a third task where difficulty is manipulated differently may have had an indirect effect on participants’ disposition towards their confidence ratings in the detection and discrimination tasks. An alternative interpretation is that in our previous study a difference between detection and discrimination was a false positive driven by noise, which happened to facilitate the quadratic trend in one task and reduce it in another. Indeed, this difference in a quadratic modulation was not a-priori expected based on theory in our previous study, and was the result of exploratory analysis that went beyond our pre-registration. The current preregistered, hypothesis-driven replication provides an unbiased test of this effect, which resulted in a failure to replicate it.

Surprisingly, however, the third ‘hybrid’ tilt-recognition task which we introduced - a discrimination task with the signal detection properties of a detection task - produced the strongest quadratic modulation of confidence in a number of regions of interest. Specifically, in BA46, rTPJ and rSTS, a quadratic modulation of confidence was significantly more pronounced in tilt recognition than in the discrimination task. Although this result is difficult to interpret without it being accompanied by a significant difference in quadratic modulation for detection (and thereby being consistent with a signature of unequal variance), it nevertheless provides some support that a quadratic modulation of confidence-related brain activity may be sensitive to the variance structure of perceptual evidence. Indeed, the hybrid tilt-recognition task showed exaggerated behavioural markers of unequal variance compared to both the detection and discrimination tasks (in both zROC curves and rcROCs; see Fig. 2). Our results are therefore consistent with a graded sensitivity of this neural marker to variance structure. In contrast, the fact that the strongest quadratic modulation was observed in decisions about stimulus category (tilt vs. vertical), rather than stimulus presence or absence, strongly suggests that this effect is not driven by a qualitative difference between detection and discrimination decisions..

### A similar linear modulation of confidence for detection ‘yes’ and ‘no’ responses in rTPJ

In our previous study, whole-brain analysis revealed a cluster in the right posterior TPJ in which a linear modulation of decision confidence was more negative for detection ‘no’ compared to ‘yes’ responses. We interpreted this finding in light of a potential role for attention monitoring in inference about absence, where participants are required to differentiate failures to perceive the target due to target absence and due to lapses of attention, and in light of an involvement of the temporoparietal junction in modelling attention states (Graziano & Webb, 2015).

In this second experiment, we pre-registered our plan to directly test this hypothesis, defining the rTPJ based on the relevant contrast in our first study. Contrary to our expectation, a negative modulation of decision confidence in the rTPJ ROI was similar for the two detection responses (see S13). A Bayesian t-test provided moderate evidence for the null hypothesis that confidence encoding in this region is invariant to detection response.

### Multivariate analysis provides weak evidence for distinct confidence encoding in an unequal-variance setting

Using Representational Similarity Analysis, we compared multivariate activation patterns in our pre-specified ROIs against theory-driven representational similarity matrices (see Fig. 4). This analysis revealed robust encoding of decision confidence in BA46 and vmPFC, consistent with a representation of decision confidence that is either task invariant or sensitive to the variance structure of a task. In contrast, multivariate activation patterns in rSTS were most consistent with a representation of detection decisions about stimulus presence or absence. Follow-up multiple regression analysis revealed that rSTS similarly encoded decision confidence in the two discrimination tasks, but that these confidence representations were not shared with the detection task. This exploratory analysis provides some indirect support for different neuronal mechanisms underlying metacognitive evaluations of decisions about presence and absence, versus stimulus category.

### Power considerations in neuroimaging research

We note that our target sample size (N=35 included subjects) was based on an a-priori power calculation to obtain 65-85% power to replicate our previous findings, assuming no inflation of effect sizes in the original report, and no effect of a reduction in the number of trials per task for group-level sensitivity. Our power and sample size aspirations were balanced with resource constraints (grant funding and time, given that N=46 subjects were tested to obtain N=35 included subjects), but remain substantially above recent estimates in fMRI research, where power can be routinely lower than 10% (Cremers et al., 2017; though we note that power calculations for whole-brain analyses are complicated by the need to deal with mass-univariate multiple comparisons correction). Low statistical power hinders the field’s ability to create a progressive research programme in which one set of findings builds on the other.

Even with medium-range power of ∼75%, and assuming a true effect, we should expect mixed results in a series of studies (roughly 1 in 3 null results with an alpha level of 0.05 for individual tests). As mentioned above, our original finding of a difference in detection and discrimination was the result of exploratory analysis, likely heightening the probability of a type 1 error. Our suspicion (informed by the senior author’s experience as a younger student in the field) is that our current study would, in a previous era, have languished unpublished – a casualty of the file drawer effect. We think it is critically important that such studies are now published to allow a full picture of the strength of different effects. More generally, our findings highlight the importance of allocating funding to replication studies in cognitive neuroscience, and the importance of pre-registration of hypothesis tests. With these considerations in mind, we therefore turn in the remainder of the discussion to offer some interpretations of the current findings, and highlight questions for future research.

### A quadratic modulation of confidence: ideas and speculations

The previous sections summarise the current picture of our findings, in light of our pre-registered hypotheses, and describe both the similarities and differences between the current results and those of our previous paper. A particularly strong and consistent finding across both studies was that univariate fMRI activation in prefrontal and parietal cortex is quadratically modulated by decision confidence. However, as described above, we find no clear support for or against our hypothesised variance-structure account of a quadratic modulation of decision confidence. This therefore leaves underdetermined the computational basis of such an effect. In the following, we discuss two additional candidate interpretations of a quadratic modulation of confidence.

In Mazor et al. (2020), we referred to two previous reports of a quadratic relation between subjective ratings and brain activation: one in subjective visibility ratings (Christensen et al., 2006) and the other in product desirability ratings (De Martino et al., 2017). We then explained that our findings were qualitatively different: while a quadratic effect for visibility or product desirability can reflect a linear modulation of subjective confidence when both ends of the rating scale are associated with higher levels of confidence (for example, being highly confident that a product is or is not desirable, or that a stimulus is or is not visible), our results highlight a quadratic effect in confidence itself. In other words, the low end of the scale should reflect low confidence in the perceptual decision, rather than high confidence in a negative rating.

However, although the low end of the confidence scale reflects low confidence *in a decision*, it may still reflect a high level of confidence *in the confidence rating itself*^1^. With our incentive structure, reward is dependent not only on task performance but also on the adequacy of confidence reports (see Methods). It is therefore not unlikely a participant would reason something along the lines of “I’m not sure what I just saw, so I’m highly confident that I should rate my subjective confidence as low to maximize my bonus”. Brain regions where activation scales with subjective confidence may then reflect this meta-level subjective confidence (effectively, confidence in confidence) with a higher level of activity not only for the upper end, but also for the lower end of the confidence scale. Note this scheme does not imply the existence of ‘meta-meta-cognition’: all that is required is that confidence ratings are represented as part of the primary (type-1) task, such that metacognitive resources are available to evaluate the quality of confidence reports themselves (Lebreton, 2015).

It then remains to be explained why a quadratic effect of confidence is significantly stronger in the tilt-recognition compared to both discrimination and detection tasks (see Fig. 3 and S7). One possibility is that confidence in the presence or absence of a tilt more naturally lends itself to rule-based heuristics (such as a mapping between perceived angle and confidence level), leaving metacognitive resources free to monitor the quality of subjective confidence ratings.

Alternatively, a quadratic modulation of decision confidence may reflect object-level (that is, not meta-level) inter-trial fluctuations in visual attention. As an example of how this might arise, a recent EEG study (Davidson et al., 2021) revealed a negative linear relation between reported attention and pre-stimulus alpha power in the 8-12 HZ frequency band. In contrast, high levels of decision confidence were associated with intermediate prestimulus alpha power, giving rise to a negative quadratic relation between alpha power and confidence. One implication of this result is that brain regions where activation is typically negatively correlated with alpha power may show a positive quadratic modulation of confidence in virtue of their relation to pre-stimulus alpha. This potentially explains why a quadratic modulation of confidence is prominent in a frontoparietal network where activity has been negatively linked to alpha power (Goldman et al., 2002; Laufs et al., 2003).

It is unclear, however, why a putative relationship between neural correlates of attention and subjective confidence should be sensitive to the variance structure of the task. One useful approach is to ask how, under Bayesian decision theory, decisions and confidence estimates should adjust to reflect differences in the effects of attention on internal distributions. For instance, in a behavioural study (Denison et al., 2018), participants rated their confidence in whether the orientation of a grating was sampled from a wide or narrow distribution, both centred at 0 degrees. A comparison of trials with valid, invalid and neutral cues revealed that participants rationally adapted their decisions and confidence estimates to their current attention state. Note that this effect of attention on perceptual decisions is specific to unequal variance settings; in an equal variance setting, accuracy is highest when the decision criterion is set to midway between the two stimulus distributions, regardless of sensory precision. In contrast, an unequal variance setting introduces a link between optimal placement of the decision criterion and sensory precision. As we show in Mazor et al. (2020), a model where subjects dynamically adjust their decision criterion based on previous samples displays different associations between criterion adjustment and confidence, depending on the variance structure of the task. Together, a possible interpretation of our findings is that an quadratic modulation of confidence n regions such as BA46, rTPJ and rSTS is effectively mediated by decisions about policy changes, such as adjustments of a decision criterion based on observed samples.

### Conclusions

In conclusion, in a pre-registered experiment we find that a quadratic effect of decision confidence on brain activity is not specific to decisions about presence and absence, but may be sensitive to the variance structure of the task. We discuss three candidate accounts of this effect, one postulating a role for subjective confidence in the accuracy of confidence ratings themselves, one identifying this quadratic effect with the neural correlates of fluctuations in attention, and one linking brain activations to the online adjustment of a decision criterion.

## Methods

We report how we determined our sample size, all data exclusions (if any), all manipulations, and all measures in the study. All design and analysis details were pre-registered before data acquisition and time-locked using pre-RNG randomization: we used the SHA256 hash function to translate our <u>pre-registered protocol folder</u> to a series of bits (7c2c27da12b6768b1789907ba5d2ec46b45d302199d5368795879fcff844d043). These bits were then used to initialize Matlab’s pseudorandom number generator for determining the order and timing of experimental events (<u>relevant lines in the experimental code</u>). Doing so ensures that pre-registration could not have taken place after data collection (Mazor et al., 2019).

### Participants

46 participants took part in the study (ages 20-39, mean = 24.2 ± 4.5; 29 females). 35 participants met our pre-specified inclusion criteria (ages 20–38, mean = 25.2 ± 4.5; 24 females). We pre-specified a sample-size of 35, balancing statistical power and resource considerations. We calculated that with 35 participants, we will have statistical power of 65-85% to replicate our previous findings, assuming no inflation of effect sizes in the original report, and no effect of a reduction in the number of trials per task for group-level sensitivity.

### Design and Procedure

After a temporally jittered rest period of 500–4000 milliseconds, each trial started with a fixation cross (500 milliseconds), followed by a presentation of a target for 500 milliseconds. In all three conditions, stimuli were dynamic noisy patterns of grayscale values, out of which a grating sometimes emerged and quickly disappeared (see Fig. 1, upper panel). We chose this mode of stimulus presentation in light of indications that it produces stronger metacognitive asymmetries between the perceptions of presence and absence relative to standard static presentation modes (Maniscalco, Brain & Lau, Hakwan, 2011). Stimuli consisted of 10 grayscale frames presented at 20 frames per second within a circle of diameter 3°. Stimuli were generated in the following way:

1. Generate 10 grayscale frames (*F*_1_, …*F*_10_), each an array of 142 by 142 random luminance values.
2. Create a 142 by 142 sinusudial grating (*G*; 24 pixels per period, random phase). The orientation of the grating is determined according to the trial type.
3. Determine grating visibility for frame *i* as *p_i_* = *v* × *exp* (−|*i* − 5|/2) with being the visibility level in this trial (0 for target-absent trials).
4. For each pixel in the frame *F_i_,j,k*, replace the luminance value for this pixel with the luminance value of this pixel in the grating (*G_j_, k*) with a probability of *p_i_*.

Participants performed the following three tasks:

1. **Discrimination**. Decide whether the grating was tilted clockwise (50% of trials; 45° relative to a vertical baseline) or anticlockwise (−45° relative to a vertical baseline).
2. **Tilt Recognition**. Decide whether the grating was vertical (50% of trials; 0°) or tilted (sampled from a normal distribution with mean 0° and standard deviation). Stimuli were presented with a fixed value of 0.2 at which stimuli are clearly visible.
3. **Detection**. Decide whether the grating was present (50% of trials) or absent. Gratings in the ‘present’ trials were sampled from a normal distribution with mean 0° and standard deviation (yoked to the tilt recognition task).

For all three tasks, responses were made with the right-hand index and middle fingers, and response mappings between fingers and stimulus classes were counterbalanced between blocks.

Immediately after making a decision, participants rated their confidence on a 6-point scale by using two keys to increase and decrease their reported confidence level with their left-hand thumb, using the same procedure and incentive structure as in Mazor et al. (2020). The perceptual decision and the confidence rating phases were restricted to 1000 and 2500 milliseconds, respectively. No feedback was delivered to subjects about their performance.

Participants were acquainted with the task in a preceding behavioural session. During this session, task difficulty was adjusted independently for detection, discrimination, and tilt recognition, targeting around 70% accuracy on all three tasks. In detection and discrimination, we achieved this by adaptively controlling the visibility value once in every 10 trials: increasing it when accuracy fell below 60%, and decreasing it when accuracy exceeded 80%. In tilt recognition, was set to 0.20 such that stimuli were highly visible, and calibration was performed on the standard deviation of orientations σ_*orientation*_ in a similar manner. Performance on all three tasks was further calibrated to the scanner environment at the beginning of the scanning session, during the acquisition of anatomical (MP-RAGE and fieldmap) images. After completing the calibration phase, participants underwent five to six ten-minute functional scanner runs, each comprising one block of 26 trials from each experimental condition, presented in a random order.

To avoid stimulus-driven fluctuations in confidence, and were kept fixed within each experimental block. Nevertheless, following experimental blocks with markedly bad (≤ 52.5%) or good (≥ 85%) accuracy, *v* or σ_*orientation*_ were adjusted for the next block of the same task (divided or multiplied by a factor of 0.95 for bad and good performance, respectively).

### Scanning parameters

Scanning took place at the Wellcome Centre for Human Neuroimaging, London, using a 3 Tesla Siemens Prisma MRI scanner with a 64-channel head coil. We used the same sequences as in Mazor et al. (2020).

### Analysis

The pre-registered objectives of this study were to:

1. Replicate our finding of an interaction between task (discrimination/detection) and a quadratic effect of confidence on BOLD signal in medial and lateral frontopolar cortex, as well as in the STS and pre-SMA.
2. Replicate our finding of an interaction between detection response (‘present’/’absent’) and the linear effect of confidence on activation in the right TPJ.
3. Compare quadratic effects of confidence on activations in the frontopolar cortex, the STS and the pre-SMA in a tilt-recognition task with those in detection and discrimination tasks.
4. Compare response-specific linear effects of confidence on activation in the right TPJ in a tilt-recognition task with those in detection and discrimination tasks.

#### Exclusion Criteria

Individual experimental blocks were excluded in the following cases:

1. More than 20% of the trials in the block were missed.
2. Mean accuracy was lower than 60%.
3. The participant used the same response in more than 80% of the trials.
4. For a particular response, the same confidence level was reported for more than 90% of the trials.

The first trial of each block was excluded from all analyses, leaving 25 usable trials per block. Subjects were included only if after applying block-wise exclusion specified above, their data had at least three blocks for each task.

#### fMRI data preprocessing

As in Mazor et al. (2020), fMRI data preprocessing followed the procedure described in Morales et al. (2018). Preprocessing and construction of first- and second-level models used standardized pipelines and scripts available at https://github.com/metacoglab/MetaLabCore.

#### Regions of interest

In addition to an exploratory whole-brain analysis (corrected for multiple comparisons at the cluster level), our analysis focused on the following a priori regions of interest:

1. *Medial frontopolar cortex (FPm)* obtained from a previous connectivity-based parcellation (Neubert et al., 2014).
2. *Lateral frontopolar cortex (FPl)* obtained from (Neubert et al., 2014).
3. *Brodman area 46 (BA46)* obtained from (Neubert et al., 2014).
4. *Ventromedial prefrontal cortex (vmPFC)* defined as a 8-mm sphere around MNI coordinates [0,46,-7], obtained from a meta-analysis of subjective-value related activations (Bartra et al., 2013) and aligned to the cortical midline.
5. *Right temporoparietal junction (rTPJ)* defined using the contrast *confidence_No_ – confidence_Yes_* from Mazor et al. (peak voxel [54,-46, 26], see mask attached to the protocol folder at ‘ROIs/rTPJ.nii’).
6. *Right superior temporal sulcus (rSTS)* defined using the rSTS cluster from the contrast 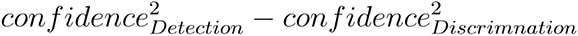 from Mazor et al. (peak voxel [60,-43,2], see mask attached to the protocol folder at ‘ROIs/rSTS.nii’).
7. *Pre-supplementary motor area (preSMA)* defined using the preSMA cluster from the contrast 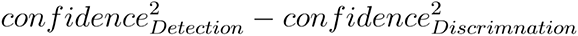 from Mazor et al. (peak voxel [0,35,47], see mask attached to the protocol folder at ‘ROIs/preSMA.nii’).

#### Univariate fMRI analysis

Univariate analysis followed a similar procedure to that described in Mazor et al. (2020). After preprocessing, runs were temporally concatenated and a design matrix fitted to the entire timecourse. Here we chose not to exclude entire runs, but specific blocks of trials. This was achieved by modelling excluded blocks with a separate nuisance regressor. We estimated two sets of design matrices:

##### Quadratic-Confidence Design Matrix (QC-DM)

The quadratic-confidence design matrix for the univariate GLM analysis consisted of 18 regressors of interest. First, a regressor for each of the six responses: ‘yes’, ‘no’, ‘tilted’, ‘vertical’, ‘clockwise’ and ‘anticlockwise’, modelled by a boxcar regressor with nonzero entries at the 4000 millisecond interval starting at the onset of the stimulus and ending immediately after the confidence rating phase, convolved with a canonical hemodynamic response function (HRF). Each of these primary regressors was accompanied by two parametric modulators, representing the linear and quadratic effects of confidence.

Trials in which the participant did not respond within the 1000 millisecond time frame, the first trial of a block, or trials in excluded blocks were modelled by separate regressors. The design matrix also included a run-wise constant term regressor, an instruction-screen regressor for the beginning of each block, motion regressors (the 6 motion parameters as extracted by SPM in the head motion correction preprocessing phase together with their first derivatives) and regressors for physiological measures (pulse and breathing). Button presses were modelled as stick functions, convolved with the canonical HRF, and separated into three regressors: two regressors for each of the two right-hand buttons, and one regressor for both up and down left-hand presses.

**Table 1.** List of regressors in the Quadratic Confidence Design Matrix (QC-DM)^2^.

|  | Task | Response |
| --- | --- | --- |
| 1 CW | Discrimination | Clockwise |
| 2 CW_conf |  |  |
| 3 CW_conf<sup>2</sup> |  |  |
| 4 ACW |  | Anticlockwise |
| 5 ACW_conf |  |  |
| 6 ACW_conf<sup>2</sup> |  |  |
| 7 Y | Detection | Yes |
| 8 Y_conf |  |  |
| 9 Y_conf<sup>2</sup> |  |  |
| 10 N |  | No |
| 11N_conf |  |  |
| 12 N_conf<sup>2</sup> |  |  |
| 13 V | Tilt recognition | Vertical |
| 14 V_conf |  |  |
| 15 V_conf<sup>2</sup> |  |  |
| 16 T |  | Tilted |
| 17 T_conf |  |  |
| 18 T_conf<sup>2</sup> |  |  |

#### Categorical-Confidence Design Matrices CC-DM

We also fitted a set of three design matrices - one for each task - in which confidence was modelled as a categorical variable. These design matrices consisted of only one regressor of interest for all included trials, modelled by a boxcar with nonzero entries at the 4000 millisecond interval starting at the onset of the stimulus and ending immediately after the confidence rating phase, convolved with a canonical hemodynamic response function (HRF). This regressor was in turn modulated by a series of 12 dummy (0/1) parametric modulators - one for every response (‘yes’ and ‘no’ for detection, ‘vertical’ and ‘tilted’ for tilt recognition and ‘clockwise’ and ‘anticlockwise’ for discrimination) and confidence rating (1-6 for both tasks). Using three design matrices instead of one allowed us to set trials from the remaining two tasks to serve as a baseline for the task of interest. These design matrices included the same set of nuisance regressors as the main design matrix.

For each participant, beta-estimates from the categorical-confidence design matrices were used as input to six response-specific multiple linear regression models, with linear confidence and quadratic confidence terms as predictors, in addition to an intercept term. Subject-specific coefficients were then subjected to ordinary least squares group-level inference, to estimate linear and quadratic effects of confidence on univariate brain activation and compare these effects between responses. The rationale for employing this two-step approach is its indifference to differences in the confidence distributions for the six responses, which may bias the estimation of quadratic and linear terms.

### Representational Similarity Analysis (RSA)

Representational Similarity Analysis (Kriegeskorte et al., 2008) was used to detect consistent spatiotemporal structures in the representation of choice and confidence across tasks and responses, within our 7 pre-specified ROIs. High and low confidence trials were defined using a median split within each response category. The empirical Representational Dissimilarity Matrix (RDM) was then compared against the following set of a-priori theoretical RDMs:

1. **Task** (Fig. 4a). Trials of the same task are similar, trials of different tasks are different.
2. **Variance structure** (Fig. 4b). Discrimination trials are similar to each other. Detection ‘present’ trials and tilt recognition ‘tilted’ trials are similar (high variance), and detection ‘absent’ trials are similar to tilt recognition ‘vertical’ trials (low variance).
3. **Decision: detection only** (Fig. 4c). Detection ‘yes’ responses are different from detection ‘no’ responses, with no consistent differences between tilt-recognition or discrimination responses.
4. **Decision: unequal variance only** (Fig. 2d). Detection ‘yes’ responses are different from detection ‘no’ responses, and ‘tilted’ responses are different from ‘vertical’ responses in the tilt recognition task, with no consistent differences between the two discrimination responses.
5. **Confidence** (Fig. 4e). High and low confidence trials are represented differently, without an effect of task or response.
6. **Confidence and variance structure interaction** (Fig. 4f). High and low confidence trials are represented differently. This effect is modulated by the variance structure of the trial category.
7. **Confidence in detection only** (Fig. 4g). High and low confidence trials are represented differently in detection only.
8. **Confidence in unequal variance only** (Fig. 4h). High and low confidence trials are represented differently in detection and tilt recognition only.

Since tasks were presented in different blocks, a high degree of similarity between conditions within a task could emerge due to temporal autocorrelations in physiological and physical noise, irrespective of distances in neural representations. To control for this, neural RDMs were constructed from distances between pairs of conditions from distinct experimental runs, and never within a single run. We chose Euclidean distance as our dissimilarity measure in order to be sensitive to differences in overall activity between tasks and responses, in addition to the relative activation patterns of voxels within an ROI.

For each ROI, the lower bound of the noise ceiling was defined as the average Spearman correlation between a given participants’ empirical RDM and the average ranked RDM of all other participants (Nili et al., 2014). This number reflects the shared variance between the RDMs of different participants that can be captured by any theoretical RDM.

### Group level inference

For exploratory whole-brain analysis, group level inference followed an ordinary least squares (OLS) procedure on the subject-specific contrast maps. Correction for multiple comparisons was performed at the cluster level, using a significance threshold of P=0.05 and a cluster defining threshold of P=0.001. No correction for multiple comparisons was applied to our prespecified ROIs.

Bayes factors were extracted by following the method described in Rouder et al. (2009). Whenever an expected effect size could be estimated from Exp. 1, we used it as a scaling factor for the prior distribution over effect sizes, reflecting a belief that if an effect exists, there is a probability of 0.5 that it is weaker than what we had observed in Exp. 1. Whenever an effect size could not be reliably estimated, we used the default scaling factor of 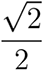.

## Supplementary information

**S1.**
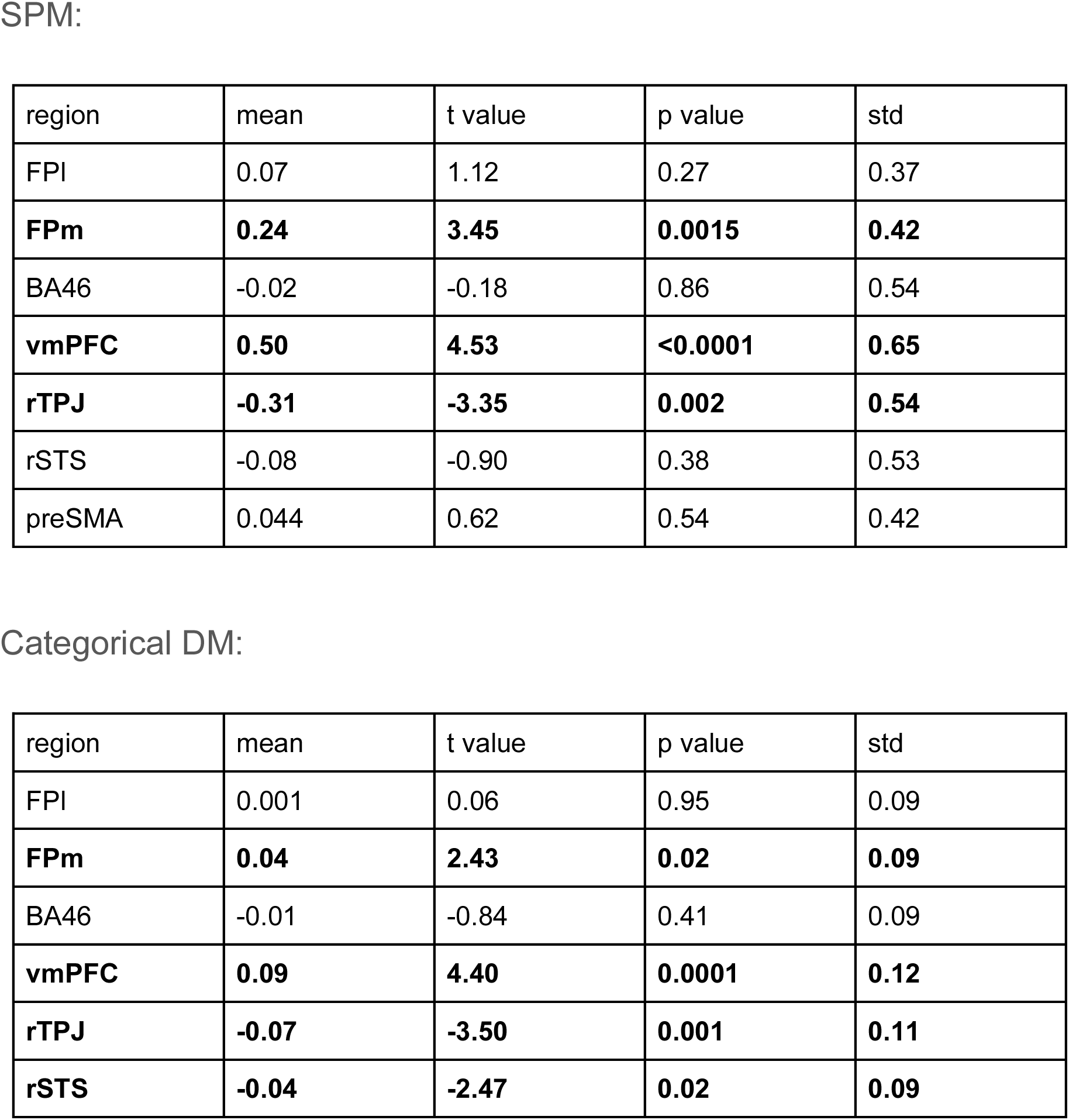
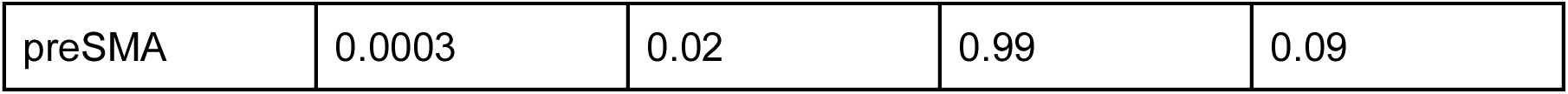
Linear effect of confidence across the seven ROIs: In S1-S8, we use the following coding to indicate significance: - **Significant at the 0.05 level (uncorrected)** - *Bayes Factor provides evidence for the null (BF_01_>3)*

**S2.**
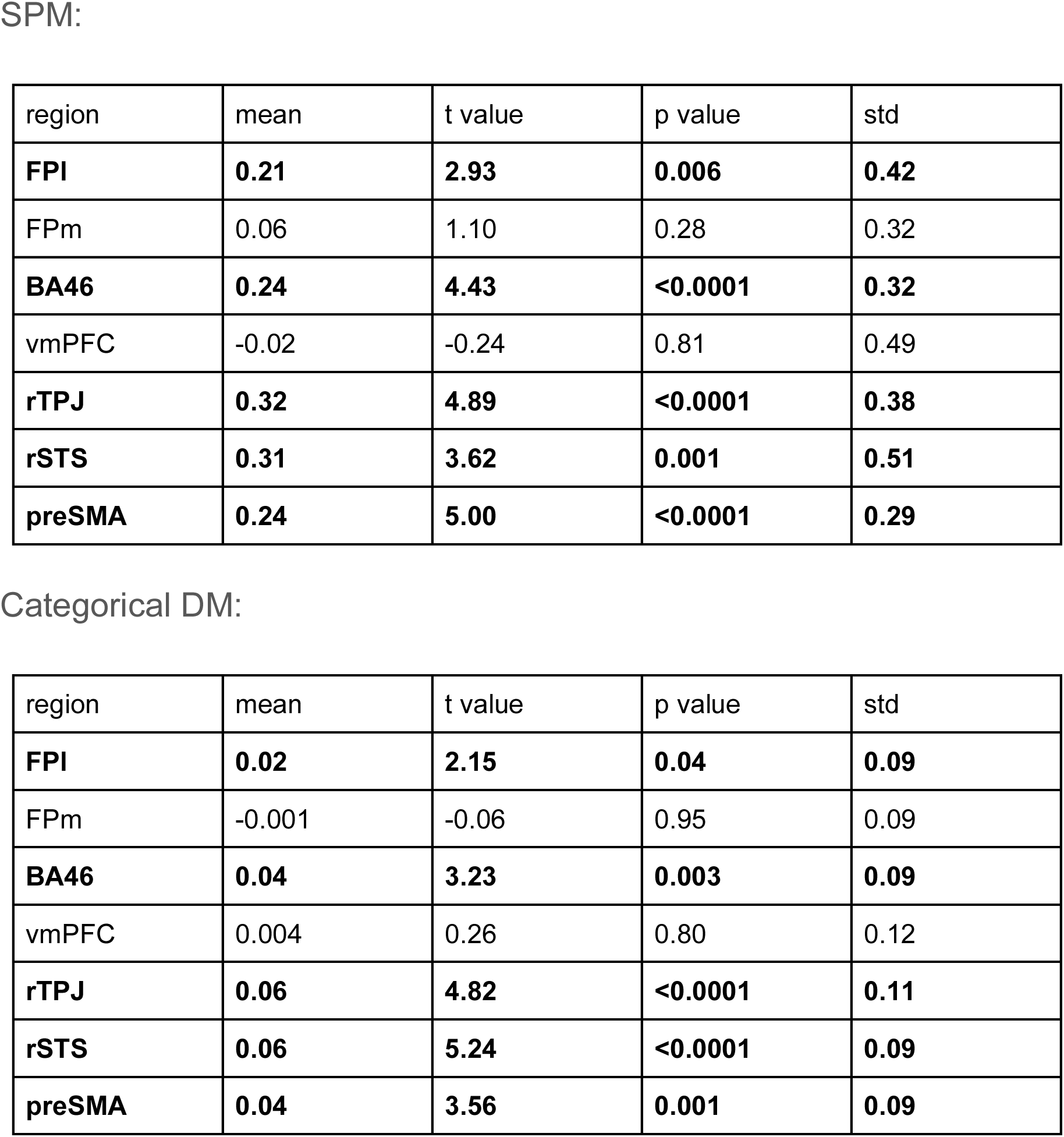
Quadratic effect of confidence across the seven ROIs:

**S3.**
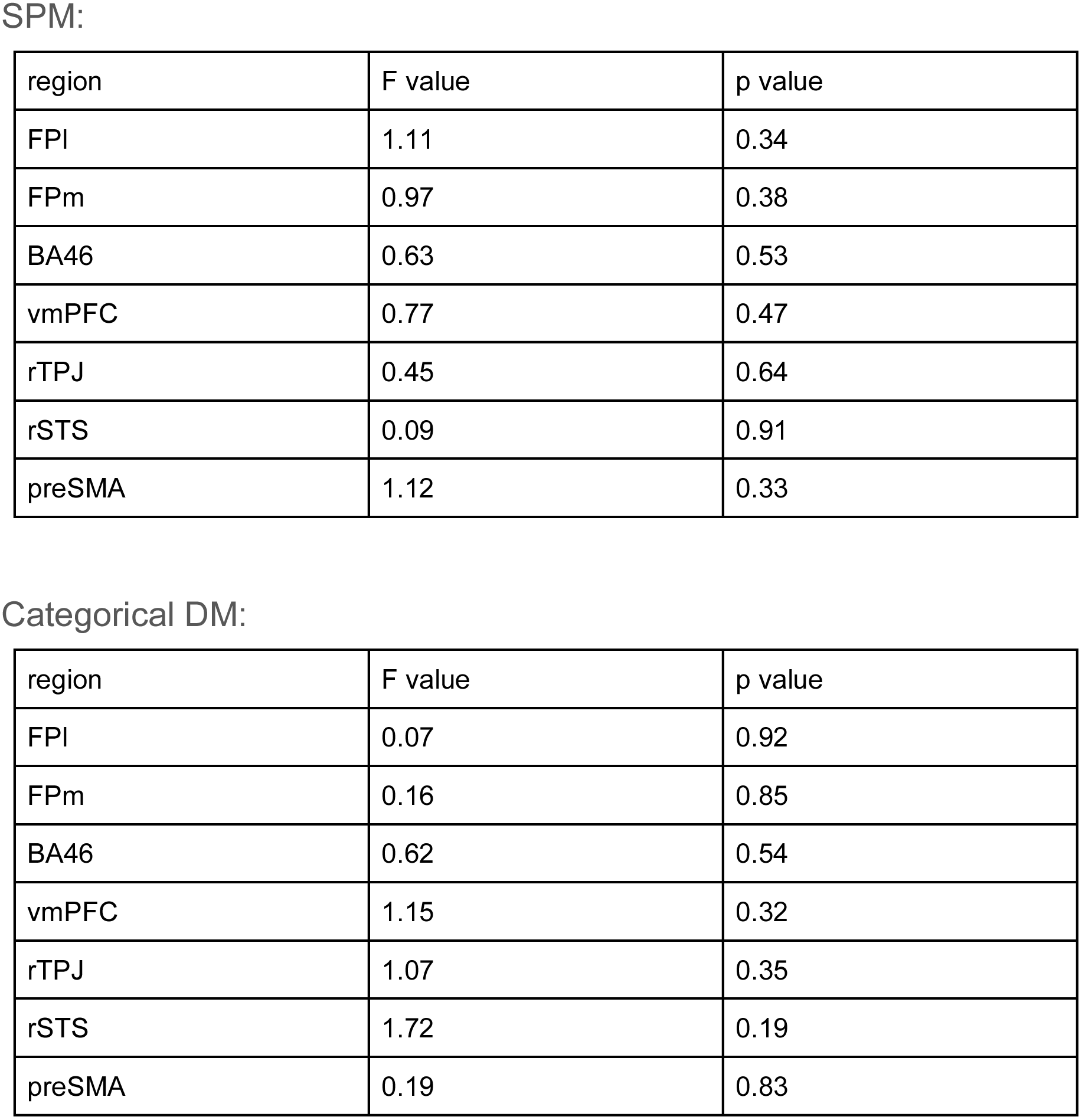
Main effect of task across the seven ROIs:

**S4.**
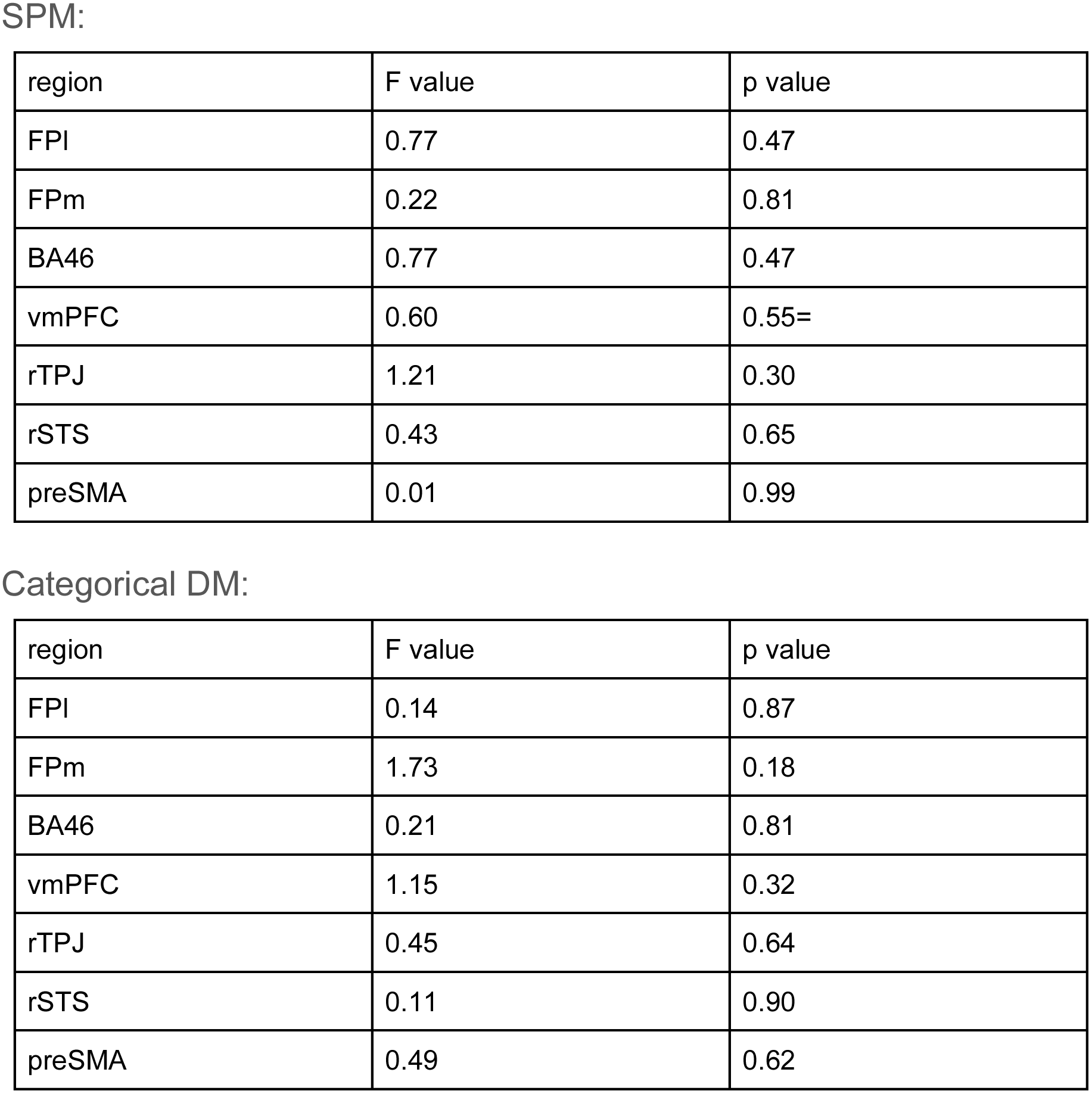
Interaction between task and linear effect of confidence across the seven ROIs:

**S5.**
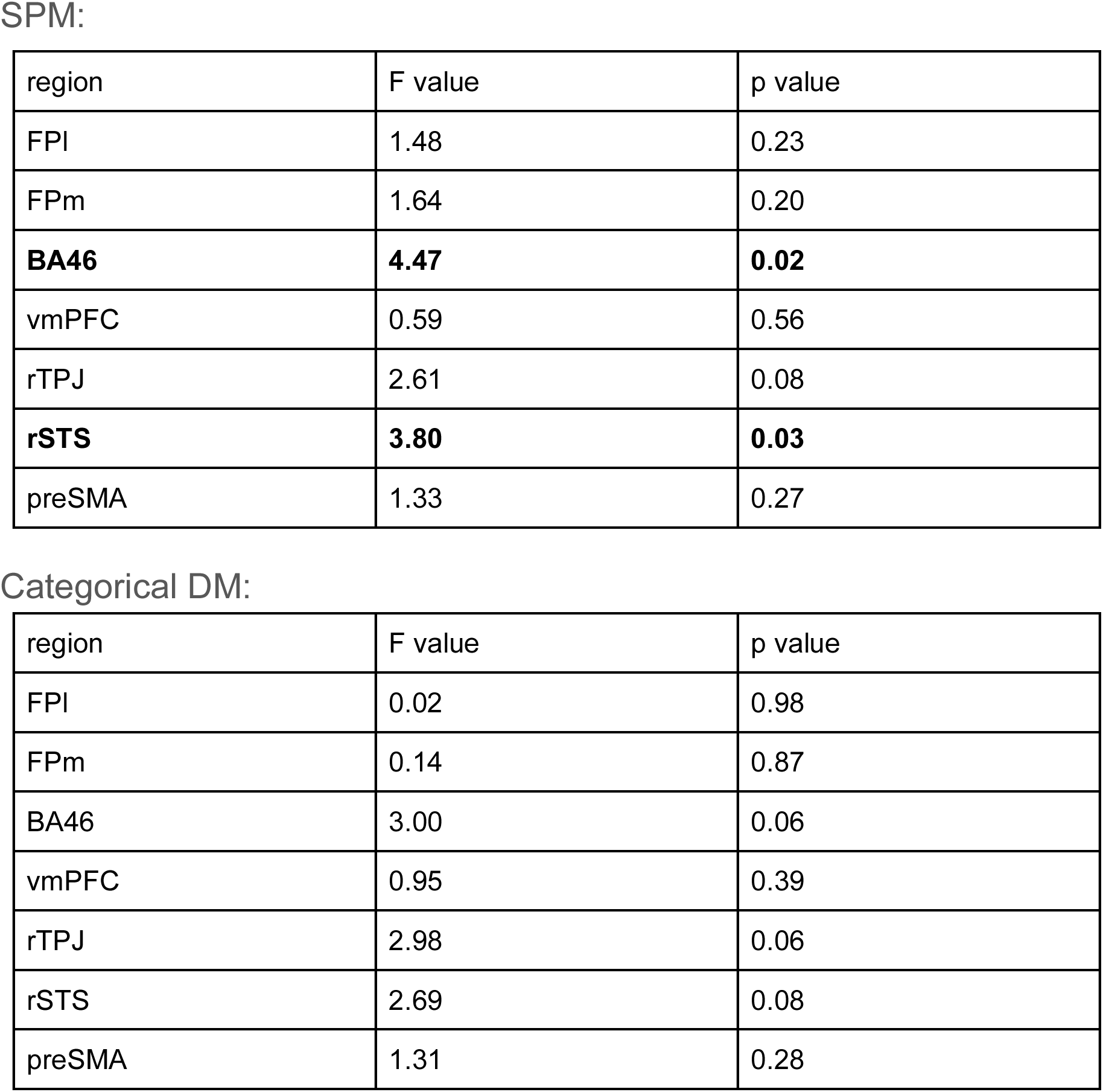
Interaction between task and quadratic effect of confidence across the seven ROIs:

**S6.**
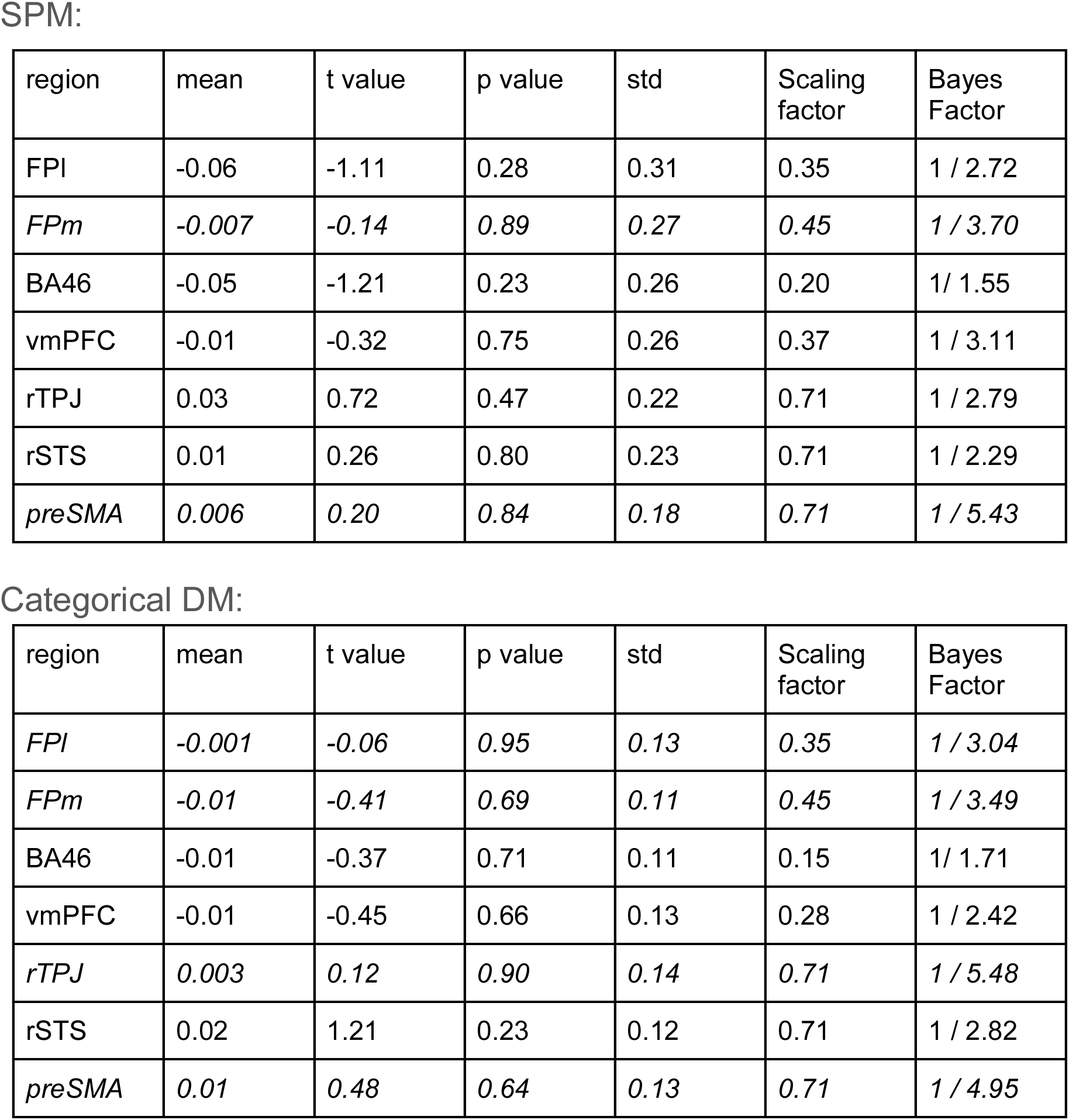
Quadratic effect of confidence in detection versus discrimination across the seven ROIs:

**S7.**
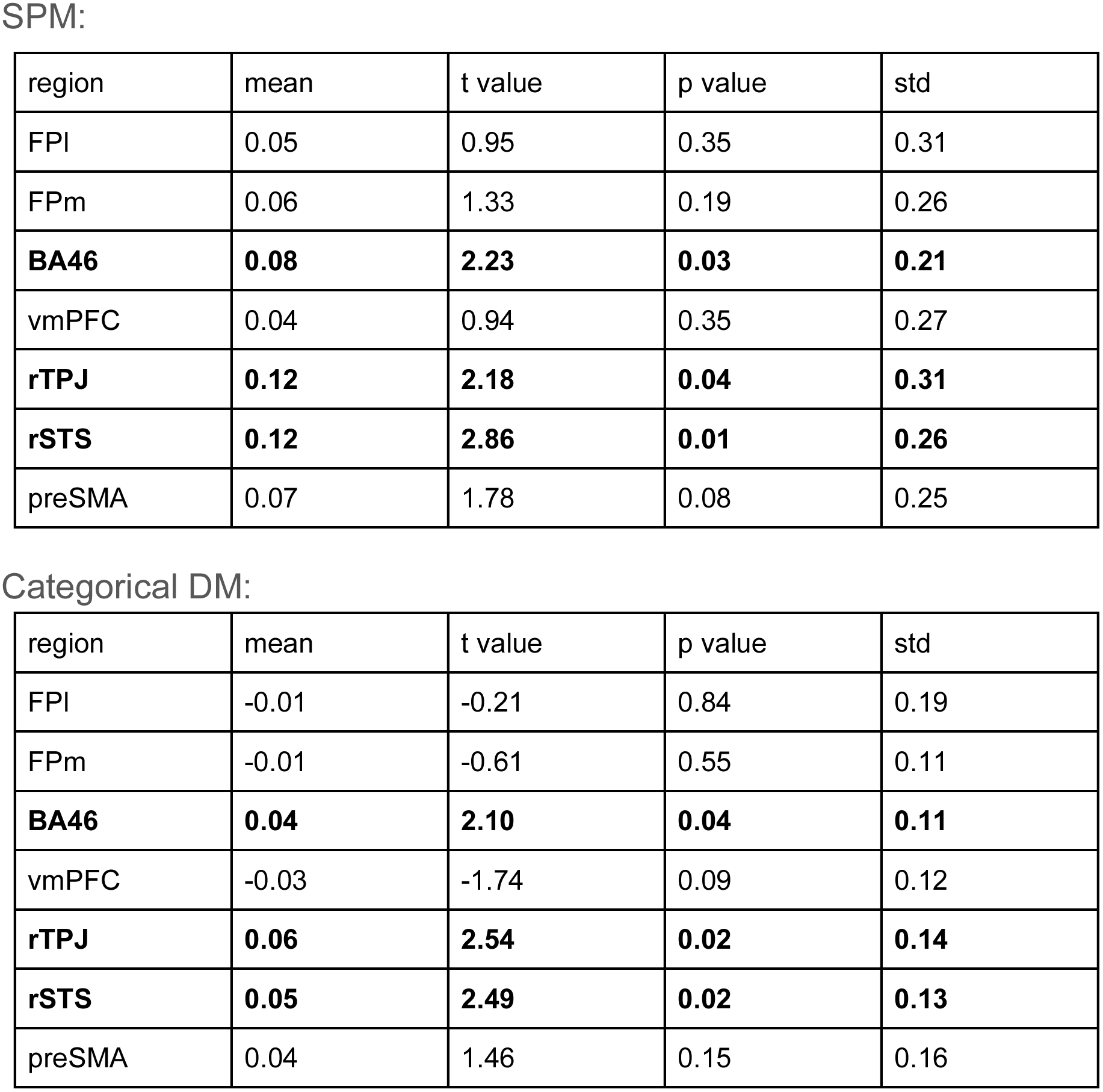
Quadratic effect of confidence in tilt recognition versus discrimination across the seven ROIs:

**S8.**
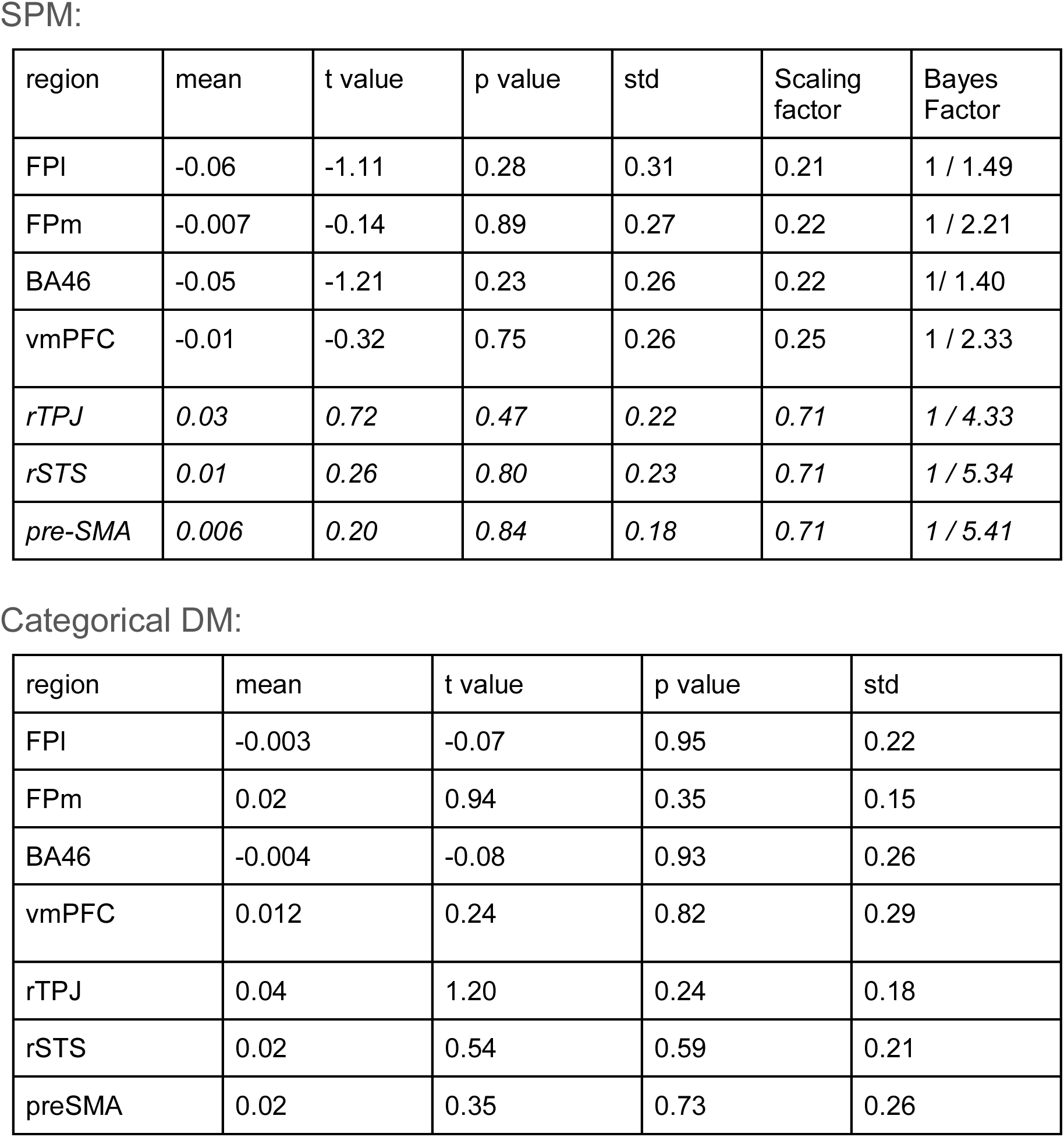
Linear effect of confidence in detection ‘yes’ versus ‘no’ responses across the seven ROIs:

**S9.**
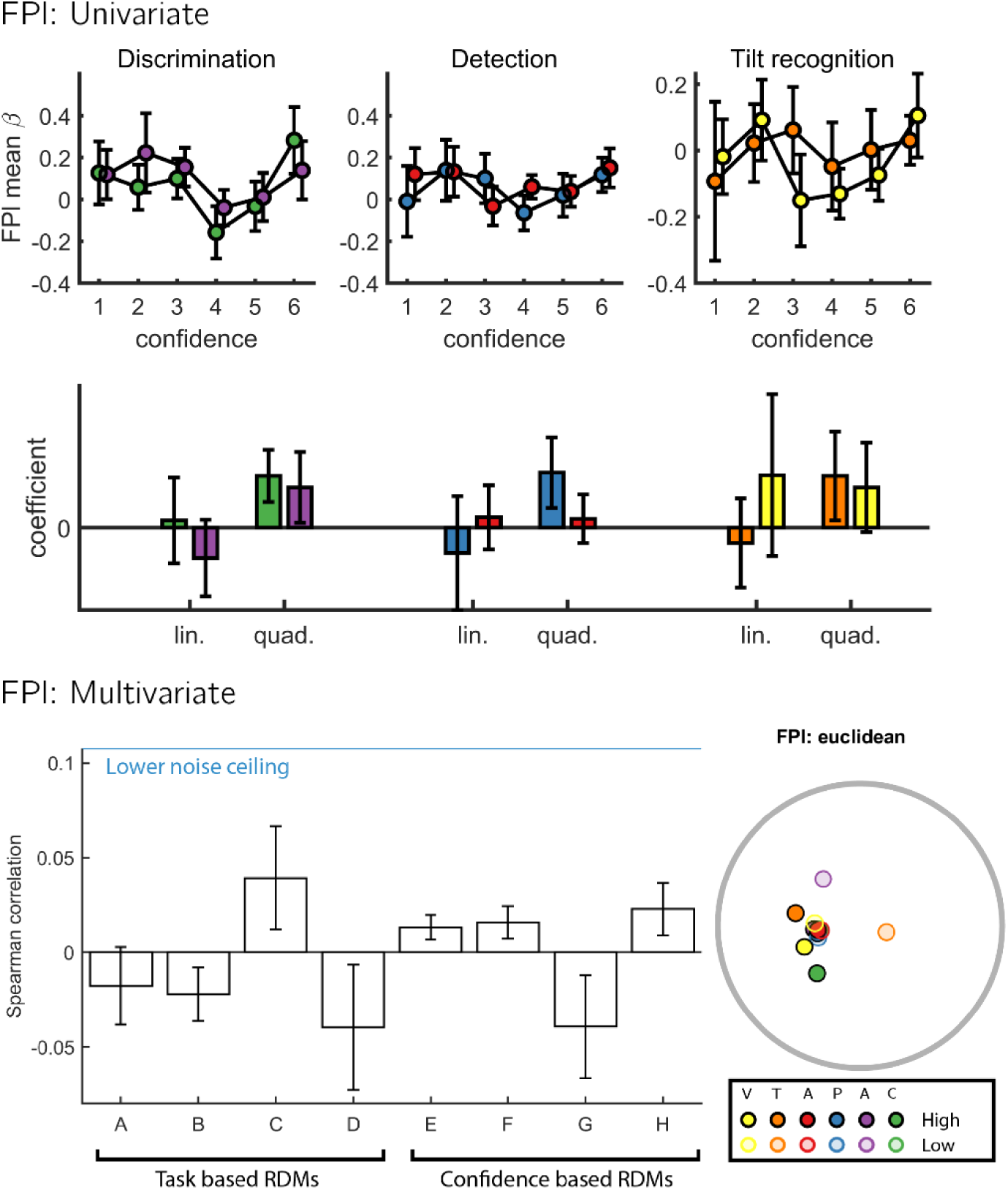
Full results: FPl. First row: mean beta coefficients per response and confidence level from the categorical design matrix. Second row: mean beta coefficients from a multiple regression model fitted to the above curves, using confidence and confidence squared as predictors. Third row: Mean Spearman correlations between the empirical RDM and theory driven RDMs. Error bars represent the standard error of the mean. Bottom right corner: A multidimensional scaling projection of the empirical RDM onto a two-dimensional space, for the purpose of visualization. In the FPl ROI, multivariate spatial activation patterns did not significantly correlate with any of the eight theoretical RDMs (p’s>0.05).

**S10.**
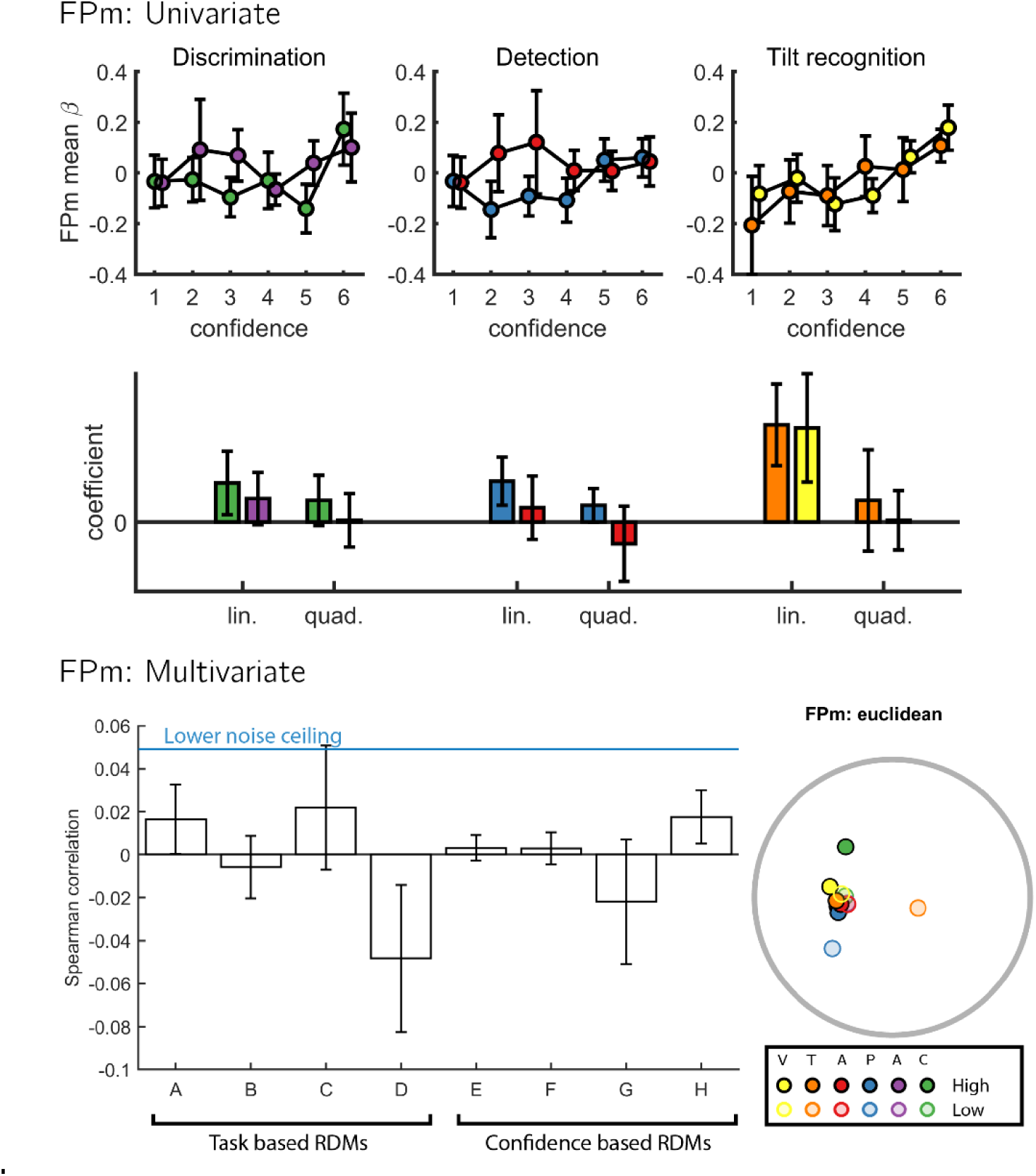
Full results: FPm. In the FPm ROI, multivariate spatial activation patterns did not significantly correlate with any of the eight theoretical RDMs (p’s>0.16). Same conventions as Fig. S9.

**S11.**
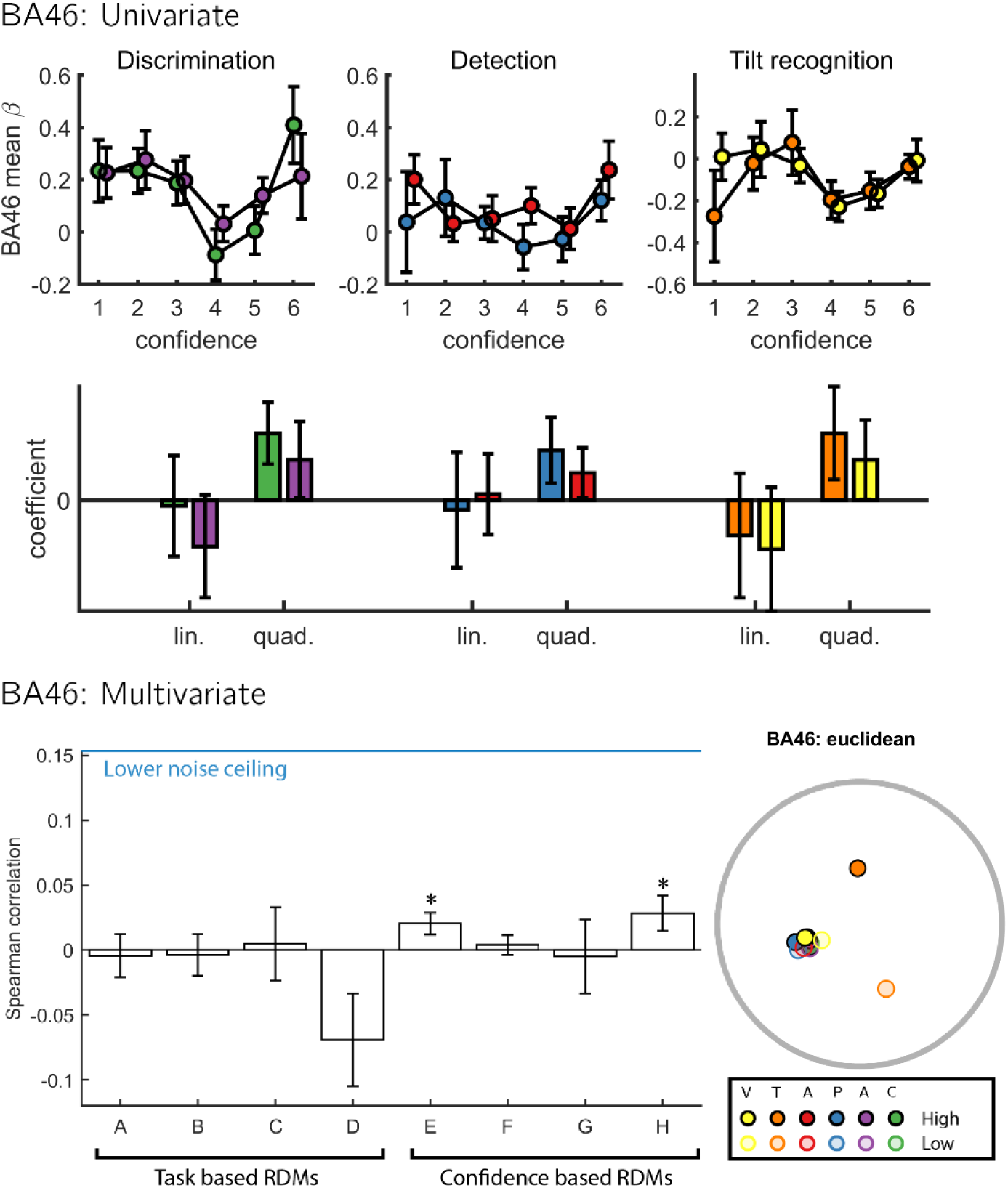
Full results: BA46. In the BA46 ROI, multivariate spatial activation patterns corresponded most with RDM E, where confidence is encoded in a task-invariant manner (*t(34)=2.38, p<0.05*), and with RDM H, where confidence is encoded in unequal variance only (*t(34)=2.08, p<0.05*). Direct comparisons revealed no difference between the two correlations (*t(34)=2.23, p<0.05*) and marginally with RDM E (*t(34)=0.60, p=0.55*). Same conventions as Fig. S9.

**S12.**
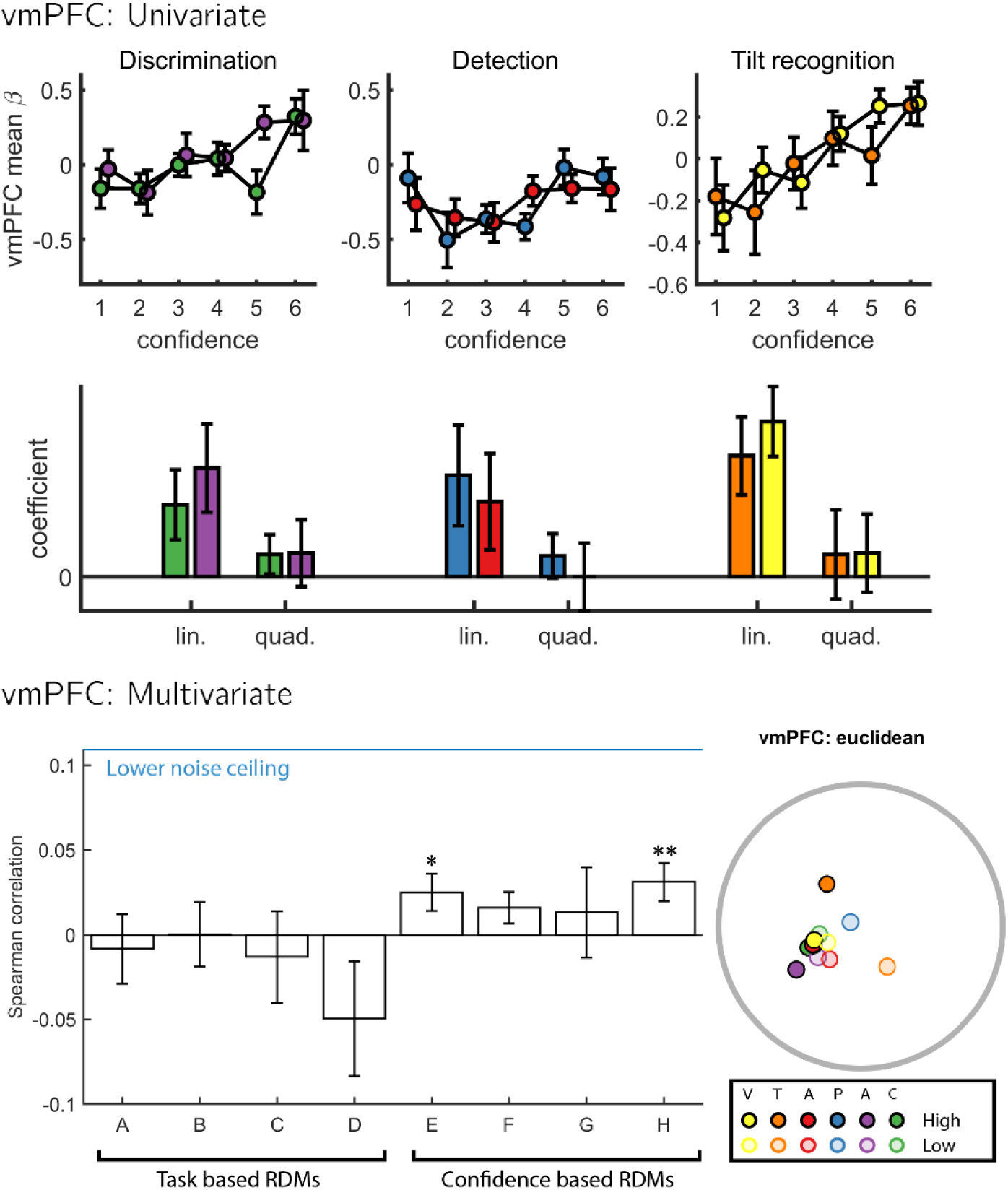
Full Results: vmPFC. In the vmPFC ROI, multivariate spatial activation patterns corresponded most with RDM H, where encoding of confidence was selective to unequal variance only (*t(34)=2.76,p<0.01*), and with RDM E, where confidence is encoded in a task-invariant manner (*t(34)=2.26, p<0.05*). Direct comparisons revealed no differences in the fits of these two RDMs to activation patterns in this region (t(34)=0.54,p=0.59). Same conventions as Fig. S9.

**S13.**
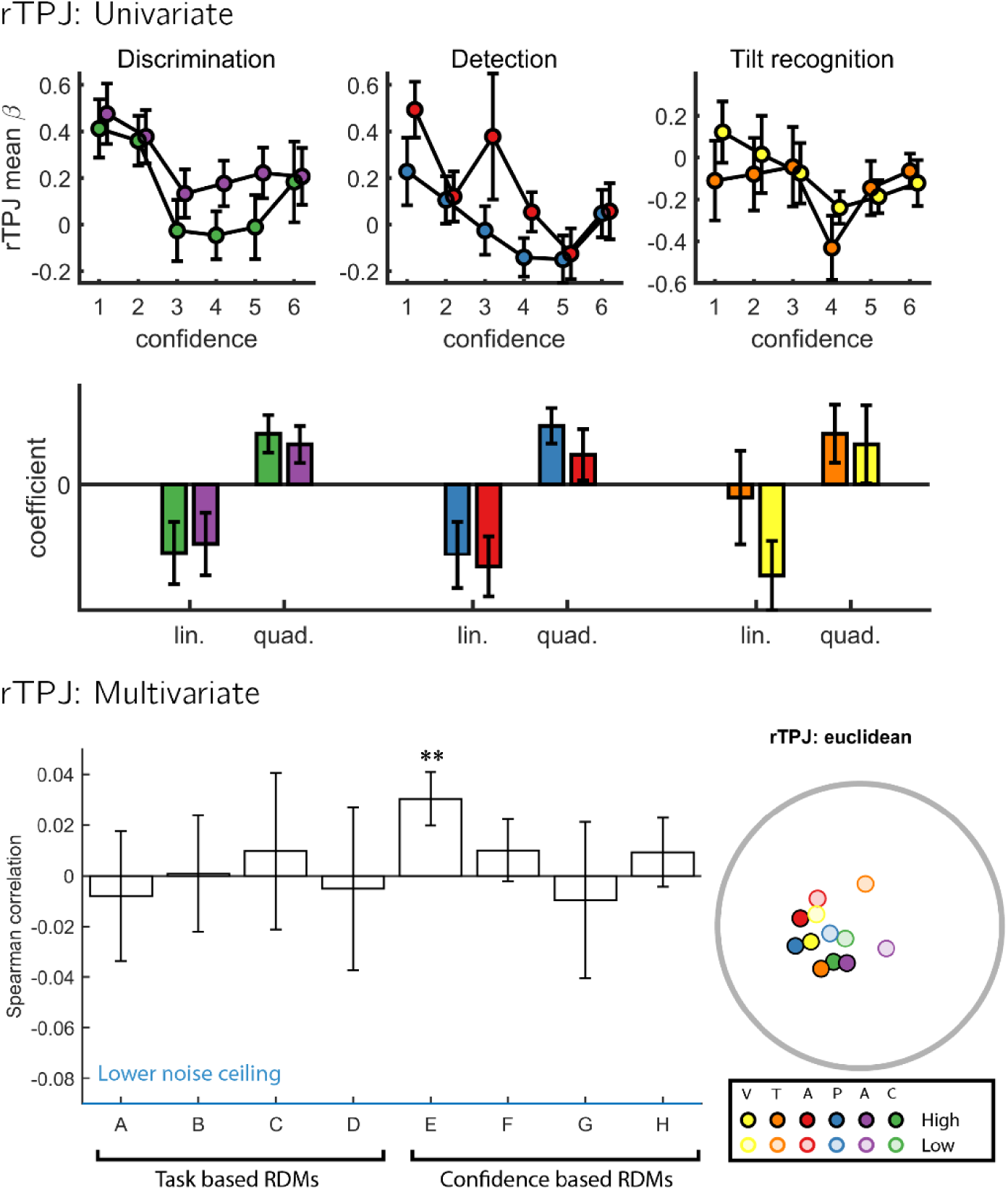
Full Results: rTPJ. In the rTPJ ROI, multivariate spatial activation patterns corresponded most with RDM E, where encoding of confidence was independent of task (*t(34)=2.86, p<0. 01*). Same conventions as Fig. S9.

**S14.**
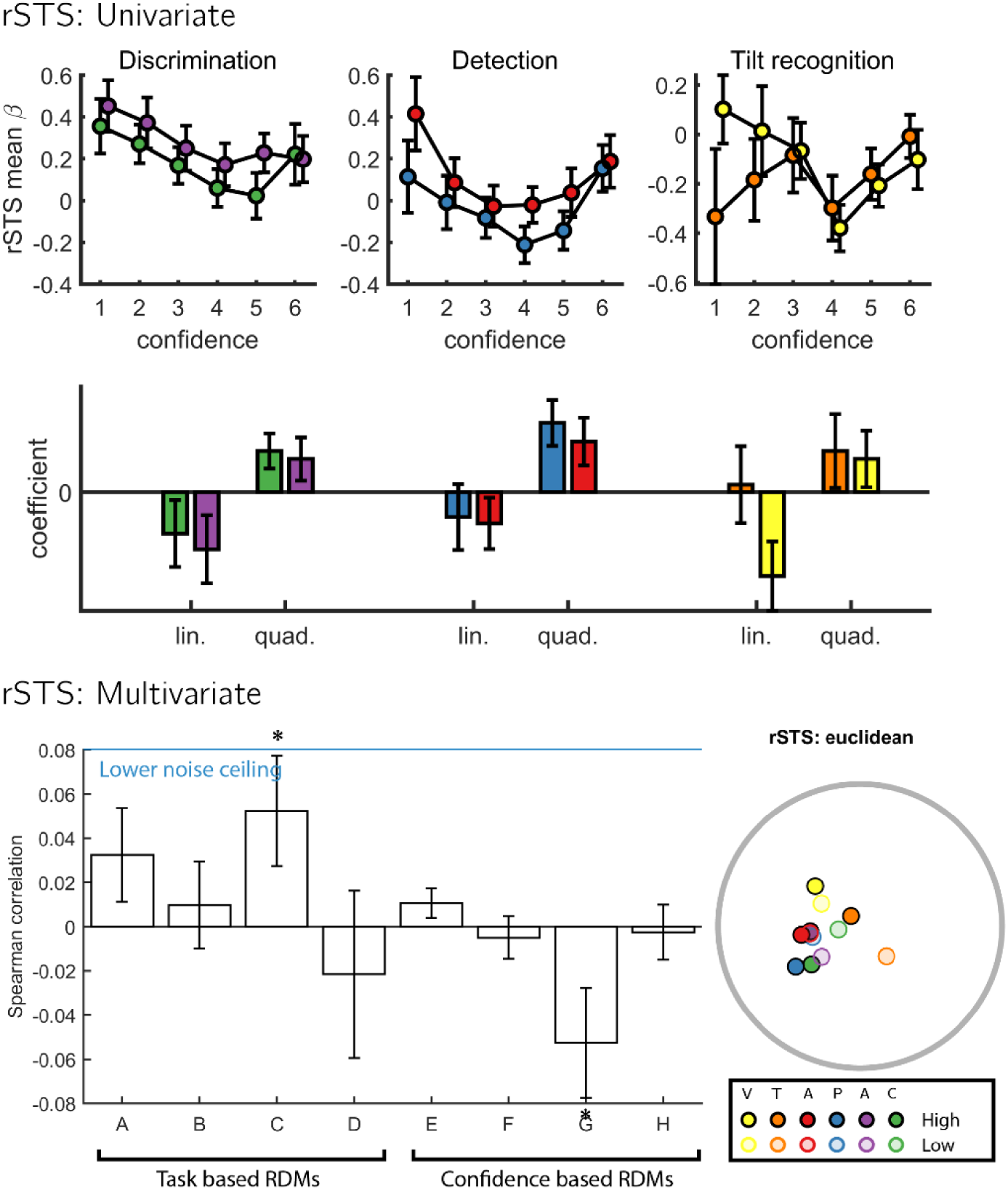
Full Results: rSTS. In the rSTS ROI, multivariate spatial activation patterns corresponded most with RDM C, where detection responses are encoded (*t(34)=2.10,p<0.05*). A negative correlation with RDM G reflected the fact that when ignoring the diagonal, RDM G is the perfect opposite of RDM C. Same conventions as Fig. S9.

**S15.**
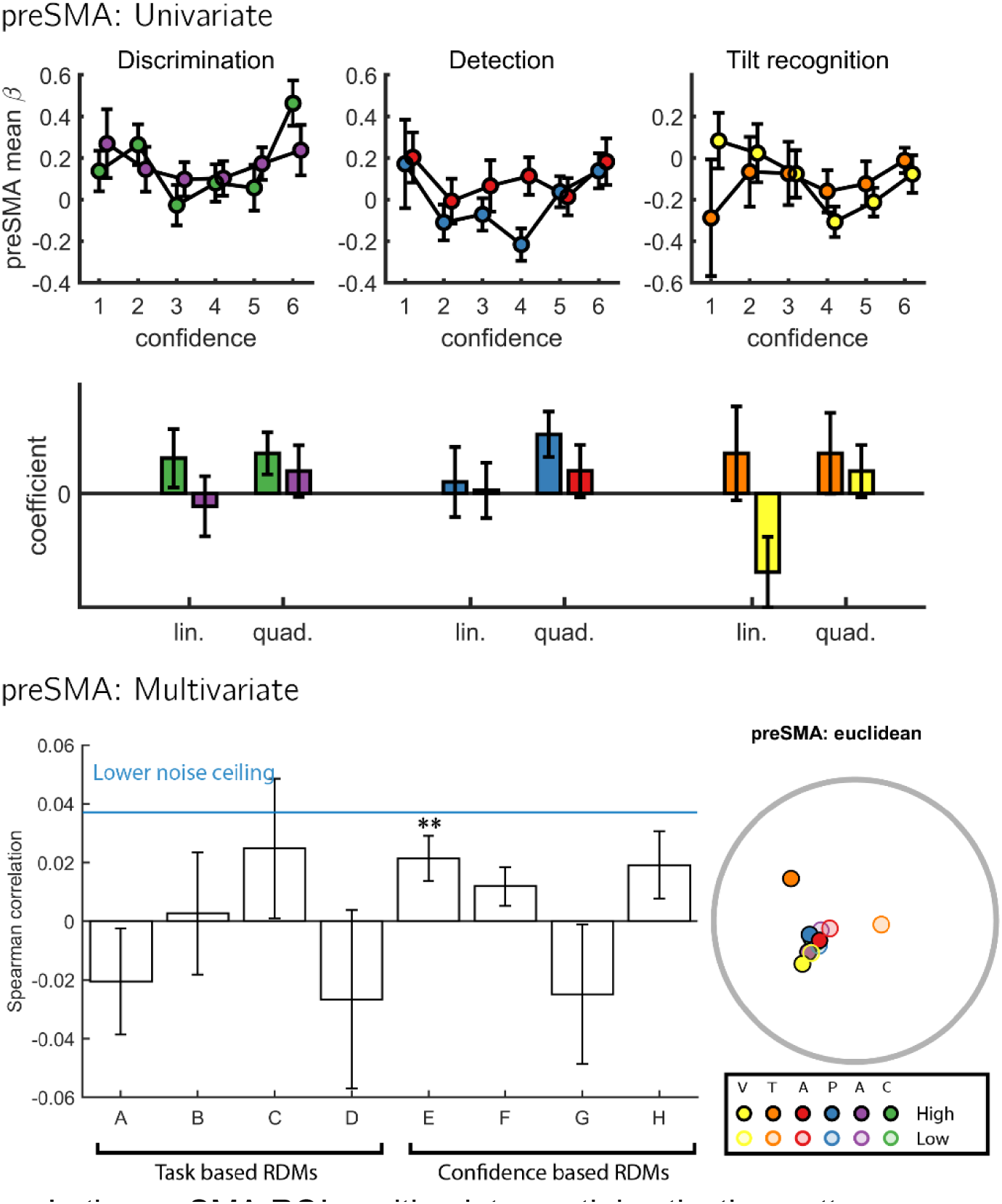
Full Results: pre-SMA. In the preSMA ROI, multivariate spatial activation patterns corresponded most with RDM E, where encoding of confidence was independent of task (*t(34)=2.76,p<0.01*). Same conventions as Fig. S9.

## Footnotes

1 We thank Heleen Slagter for suggesting this idea.

2 Due to a mistake, our pre-registration document specified that trials will be classified based on the presented stimulus rather than the response. This would mean that high confidence hits would be classified together with high confidence misses, as in both cases the stimulus was a grating. We therefore decided to follow the analysis strategy of Mazor et al. (2020), and classify trials based on response instead.

